# Human organoids reveal PTEN-driven mesendoderm specification via retinoic acid signaling suppression

**DOI:** 10.1101/2025.10.17.682808

**Authors:** Wuming Wang, Qiqi He, Hong Zhang, See-Wing Chan, Zhiqiang Xiong, Haite Tang, Yue Lv, Xianwei Su, Ya Guo, Yong Fan, Xingguo Liu, Xiuwu Bian, Andrew M. Chan, Hongbin Liu, Gang Lu, Wai-Yee Chan

## Abstract

During early embryogenesis, epiblast cells ingress through the primitive streak (PS) and commit to a mesendoderm (MES) fate, giving rise to both mesodermal and endodermal lineages. Despite advancements in human organoid technology for modeling pre-implantation blastocyst stage, robust systems that accurately recapitulate MES specification and PS formation during the post-implantation stage remain limited. Here, we show that human induced pluripotent stem cells can be used to generate three-dimensional MES organoids that faithfully mimic *in vivo* human MES specification, particularly from the anterior PS. We demonstrate that MES organoid formation is dependent on *PTEN* expression and that loss of *PTEN* severely impairs the formation of mesoderm and definitive endoderm derivatives arising from MES organoids. To elucidate the molecular mechanisms underlying these defects, we conducted an integrative multi-omics analysis, including transcriptome, chromatin accessibility profiling, metabolism, and phosphoproteomics, on wild-type and *PTEN^−/−^*MES cells. These analyses revealed that PTEN suppresses retinoic acid (RA) signaling during MES lineage commitment. Furthermore, we identified the RA-degrading enzyme CYP26A1 as a downstream effector of PTEN. Notably, while excessive RA induced by *PTEN* ablation is detrimental to MES cell generation, physiological levels of RA are necessary. Collectively, our human MES organoids offer a valuable model for dissecting early human development, and our findings identify PTEN as a key regulator of MES fate commitment through inhibition of RA signaling.

## Introduction

The formation of the primitive streak (PS)—a transient, thickened region of the epiblast— marks a pivotal event during gastrulation. Within the PS, epiblast cells acquire distinct fates in a spatially and temporally regulated manner. Cells ingressing into the anterior PS (APS) adopt a mesendoderm (MES) identity, co-expressing lineage-specifying transcription factors such as *EOMES*, *SOX17*, *GSC*, *FOXA2*, and *LHX1* ^1^. These MES cells then undergo epithelial–mesenchymal transition (EMT), leading to differentiation into the mesoderm and definitive endoderm (DE) ^2^. In contrast, cells from the posterior PS (PPS) contribute predominantly to the extraembryonic and somitic mesoderm ^3^.

Studying early human development poses considerable technical and ethical challenges. Recently, models such as blastoids and gastruloids derived from human induced pluripotent stem cells (iPSCs) have emerged as promising tools to examine early gastrulation events and PS formation ^4, 5, 6, 7^. While mesodermal organoids have been used to model somite development ^8^, there remains a lack of human organoid systems that robustly replicate MES specification, particularly from the APS.

The establishment of anterior-posterior (A-P) polarity is foundational for gastrulation, directing the formation and organization of the three germ layers. Canonical pathways— including Nodal, BMP, and WNT—are central to PS induction and MES cell formation in both embryos and iPSCs ^9, 10, 11^. However, BMP4 also promotes trophoblast differentiation, complicating lineage isolation, and both BMP4 and WNT support mouse embryonic stem cell self-renewal ^12, 13^, highlighting the complexity of PS regulation. Additionally, mechanical cues influence the formation of gastrulation-like structures and mesoderm specification ^14^.

Mouse epiblast stem cells have been shown to recapitulate APS transcriptional and functional features ^15^, and APS fate can also be induced in human iPSCs via YAP1 modulation ^16^. Epigenetically, the chromatin regulator Whsc1 facilitates both pluripotency exit and MES specification ^17^, while SMYD2 promotes MES generation by activating key lineage genes ^18^. Nonetheless, the precise mechanisms governing APS specification and PS organization remain incompletely understood.

Vitamin A, an essential micronutrient during pregnancy, exerts broad developmental effects but is teratogenic in excess, especially during the first 60 days post-conception ^19^. Its bioactive metabolite, retinoic acid (RA), regulates early embryonic patterning by instructing the posterior neuroectoderm and foregut endoderm while permitting trunk mesoderm development ^20^. *Pten* is indispensable for embryogenesis; *Pten*-null embryos exhibit lethality at E7.5 ^21^. Clinically, PTEN dysregulation has been linked to spontaneous miscarriage ^22, 23, 24^, further underscoring its crucial role in early development.

In this study, we developed a human MES organoid system that models core features of MES specification. These organoids can be further differentiated into DE and endothelial cell (EC) lineages, offering a tractable platform for studying early lineage commitment. We demonstrate that *PTEN* ablation disrupts MES specification *in vitro* and compromises embryonic development *in vivo*. Our results indicate that *PTEN* facilitates MES formation by modulating RA signaling via *CYP26A1*. These findings advance our understanding of human APS formation and suggest that defective MES commitment may contribute to the embryonic lethality observed in *Pten*-deficient models^21^, potentially offering mechanistic insight into the etiology of miscarriage.

## Results

### Generation of mesenoderm (MES) organoids from human iPSCs

WNT and TGF-β signaling are known to induce *in vitro* models of primitive streak (PS) formation using embryonic stem cells ^25^. In this study, we established a human MES organoid model by activating WNT signaling in iPSCs using CHIR99021 (**Figure 1a**). The resulting organoids developed a prominent layer of MES cells co-expressing *EOMES* and *SOX17*, closely recapitulating *in vivo* MES cell emergence during PS development (**Figure 1b**). Three-dimensional (3D) multiscale imaging further revealed a distinct MES-positive cell layer within the organoids (**Figure 1c**, **Supplementary Video** 1). The PPS, marked by *T*, *Meox1*, and *Kdr*, contributes to trunk mesoderm, whereas the APS, marked by *Sox17*, *Eomes, Foxa2*, and *Mixl1* gives rise to DE and cardiac mesoderm ^2, 26^. Consistent with an APS-like identity, our MES organoids expressed hallmark MES genes (*SOX17*, *EOMES*, *GSC*, *FOXA2*, *LHX1*) but lacked PPS-specific transcripts such as *CDX1*, *CDX2*, *EVX1*, *HOXA1*, and *MSGN1* (**Figure 1d**). These data indicate that the MES organoids closely model APS cells *in vivo*.

**Figure 1.**
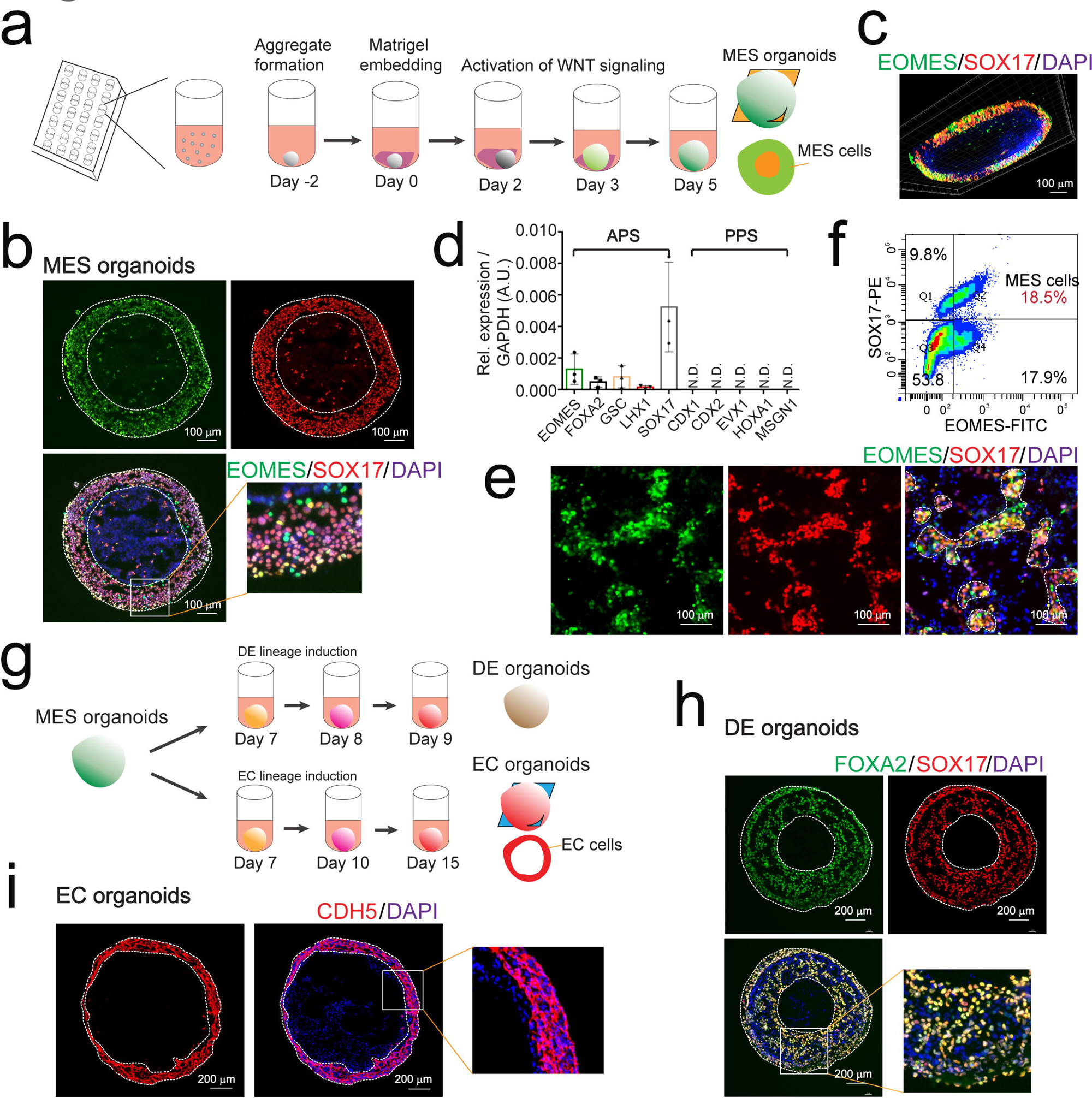
MES organoids recapitulate APS features and differentiate into DE and EC organoids. **a.** Schematic overview of the MES organoid generation protocol, wherein individual hPSC aggregates are embedded in Matrigel and differentiated via WNT pathway activation. **b.** IF staining of cryosectioned MES organoids confirming EOMES and SOX17 co-expression. Scale bar, 100 µm. **c.** Longitudinal section of a representative MES organoid showing abundant MES cells co-expressing SOX17 and EOMES. Scale bar, 100 µm. **d.** Quantitative PCR analysis of APS and PPS marker gene expression, demonstrating enrichment of APS markers. **e.** Single-cell dissociation of MES organoids followed by 24-hour re-culture to assess cellular behavior. Scale bar, 100 µm. **f.** Flow cytometric quantification of MES cells from dissociated organoids. **g.** Differentiation of MES organoids into DE and EC lineages. **h.** IF staining of DE organoid cryosections derived from MES organoids, sho--wing robust expression of DE markers SOX17 and FOXA2. Scale bar, 200 µm. **i.** EC organoid differentiation protocol and IF staining of cryosections demonstrating expression of the endothelial marker CDH5. Scale bar, 200 µm.

Temporal analysis showed significant upregulation of *EOMES* and *SOX17* expression on days 2 and 3 of differentiation (Extended Data Figure 1a, 1b). Whole-mount immunofluorescence (IF) *also* confirmed the abundance of MES cells co-expressing *EOMES* and *SOX17* within the organoids (Extended Data Figure 1c). Upon dissociation and 24-hour culture, MES-derived cells exhibited robust migratory behavior, converging into cell clusters (**Figure 1e**). Flow cytometry quantified MES cells as comprising approximately 18.5% of the organoid population (**Figure 1f**).

MES cells possess bipotentiality toward mesodermal and endodermal lineages. We leveraged this capacity by differentiating MES organoids into DE organoids, which exhibited specific expression of *SOX17* and *FOXA2* (**Figure 1g**, 1h), indicating potential for pancreatic and hepatic lineage development. To examine mesodermal potential, MES organoids were treated with VEGF and forskolin, which induced EC differentiation and the formation of EC organoids (**Figure 1g**). These were characterized by a CDH5-positive endothelial ring (**Figure 1i**). Bright-field and IF imaging revealed a chamber-like morphology (Extended Data Figure 1d), and dissociation confirmed enrichment of ECs (Extended Data Figure 1e, **Supplementary Video** 2). Together, these results highlight the developmental versatility of MES organoids and their utility in modeling early human lineage specification *in vitro*.

### Single-cell RNA-sequencing (scRNA-seq) reveals MES cell fate determination

Differentiation of iPSCs into MES organoids yields a heterogeneous population of cell types that recapitulate aspects of early embryonic development. We performed scRNA-seq to delineate the cellular composition of MES organoids. A substantial fraction of cells expressed *SOX2*, a canonical epiblast (Epi) marker (**Figure 2a**, 2c), consistent with UMAP clustering that revealed a distinct Epi population co-expressing *SOX2* and *POU5F1* (**Figure 2d**). A minor subset of cardiac progenitor cells (CPCs) was observed predominantly in the organoid periphery (**Figure 2b**), expressing markers such as *HAND1*, *HAND2*, *MYL4*, *MYL7*, and *MESP1* (**Figure 2d**).

**Figure 2.**
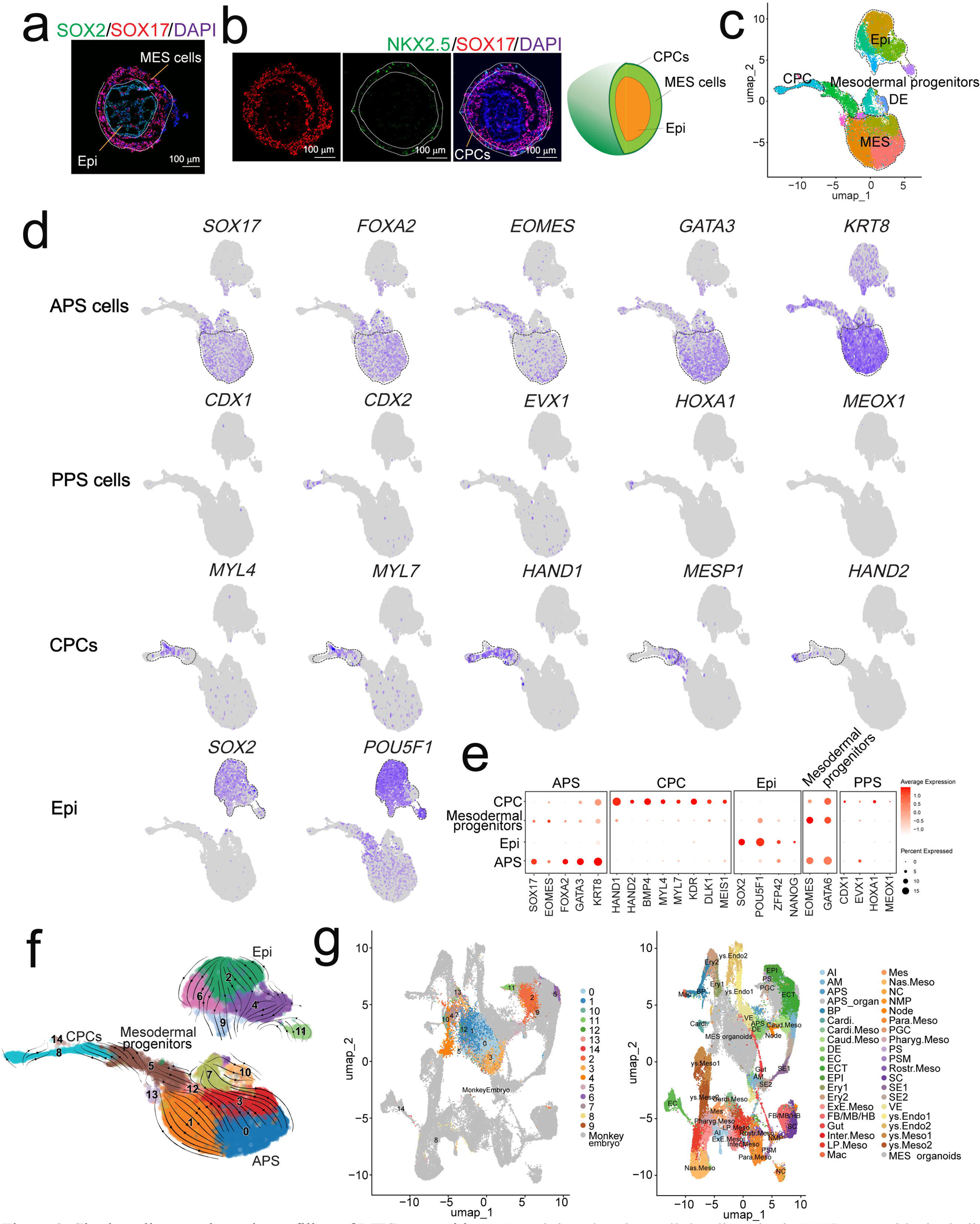
Single-cell transcriptomic profiling of MES organoids. **a.** IF staining showing cellular diversity in MES organoids, including out-layer MES cells and inner-layer Epi cells. Scale bar, 100 µm. **b.** Detection of NKX2.5-positive cells indicating the presence of CPCs on the surface of MES organoids. Scale bar, 100 µm. **c.** UMAP visualization of scRNA-seq data from MES organoids at day 3, revealing distinct cellular clusters. **d.** UMAP feature plots depicting the expression of lineage-specific marker genes corresponding to APS, PPS, CPCs, and Epi populations. **e.** Dot plot summarizing the expression profiles of lineage markers across APS, Mesodermal progenitor, Epi, and CPC clusters. **f.** RNA velocity analysis overlaid on UMAP, revealing dynamic lineage trajectories within the organoids. **g.** UMAPs showing the integration of APS organoids with reference non-human primate embryo-originated cells, demonstrating strong transcriptional similarity. Ery, erythrocyte; ys, yolk sac; Al, allan--tois; AM, amnion; FB, forebrain; Cardi., cardiac; Mes, mesenchyme; BP, blood progenitor; Mac, macrophage; ys.Endo, yolk sac endoderm.

MES cell clusters were identified based on expression of *EOMES*, *SOX17*, *FOXA2*, and *LHX1*, aligning with Q-PCR and IF staining results (**Figure 2d**; **Figure 1b–d**). These clusters lacked PPS-specific markers (*CDX1*, *CDX2*, *EVX1*, *HOXA1*, *MSGN1*), confirming their APS identity (**Figure 1d**; **Figure 2d**). A bubble plot was further used to confirm these lineage assignments (**Figure 2e**), while the violin plots show consistent marker expression across cell clusters (Extended Data Figure 2a, 2b).

RNA velocity and diffusion mapping revealed a principal developmental trajectory emerging from the Epi cluster (**Figure 2f**). To validate the identity of MES organoid-derived lineages, we compared our single-cell transcriptomes to those from monkey embryos ^27^. This integrative analysis confirmed strong transcriptional resemblance between our organoids and primate MES (APS), Epi, DE, and CPC populations (**Figure 2e**). These results validate the fidelity of MES organoids as a model for early human development and suggest their utility in uncovering mechanisms relevant to implantation failure and miscarriage.

### MES organoids reveal PTEN’s role in driving MES lineage specification

Many congenital defects and spontaneous miscarriages are driven by genetic abnormalities, including mutations in *PTEN* ^22, 23, 24, 28^. However, the role of *PTEN* during early human development remains incompletely understood. Using CRISPR/Cas9, we generated *PTEN^−/−^* iPSCs and observed that loss of *PTEN* significantly impaired MES formation. MES organoids derived from *PTEN^−/−^* cells exhibited a marked reduction in *SOX17*- and *EOMES*- positive cells (**Figure 3a**, 3b; Extended Data Figure 3a). The MES cell ring was also visibly constricted in *PTEN^−/−^* organoids (**Figure 3c**). Correspondingly, expression of key MES markers (*EOMES*, *FOXA2*, *SOX17*, *GSC*, *LHX1*) was significantly downregulated in *PTEN*- deficient cells (**Figure 3d**).

**Figure 3.**
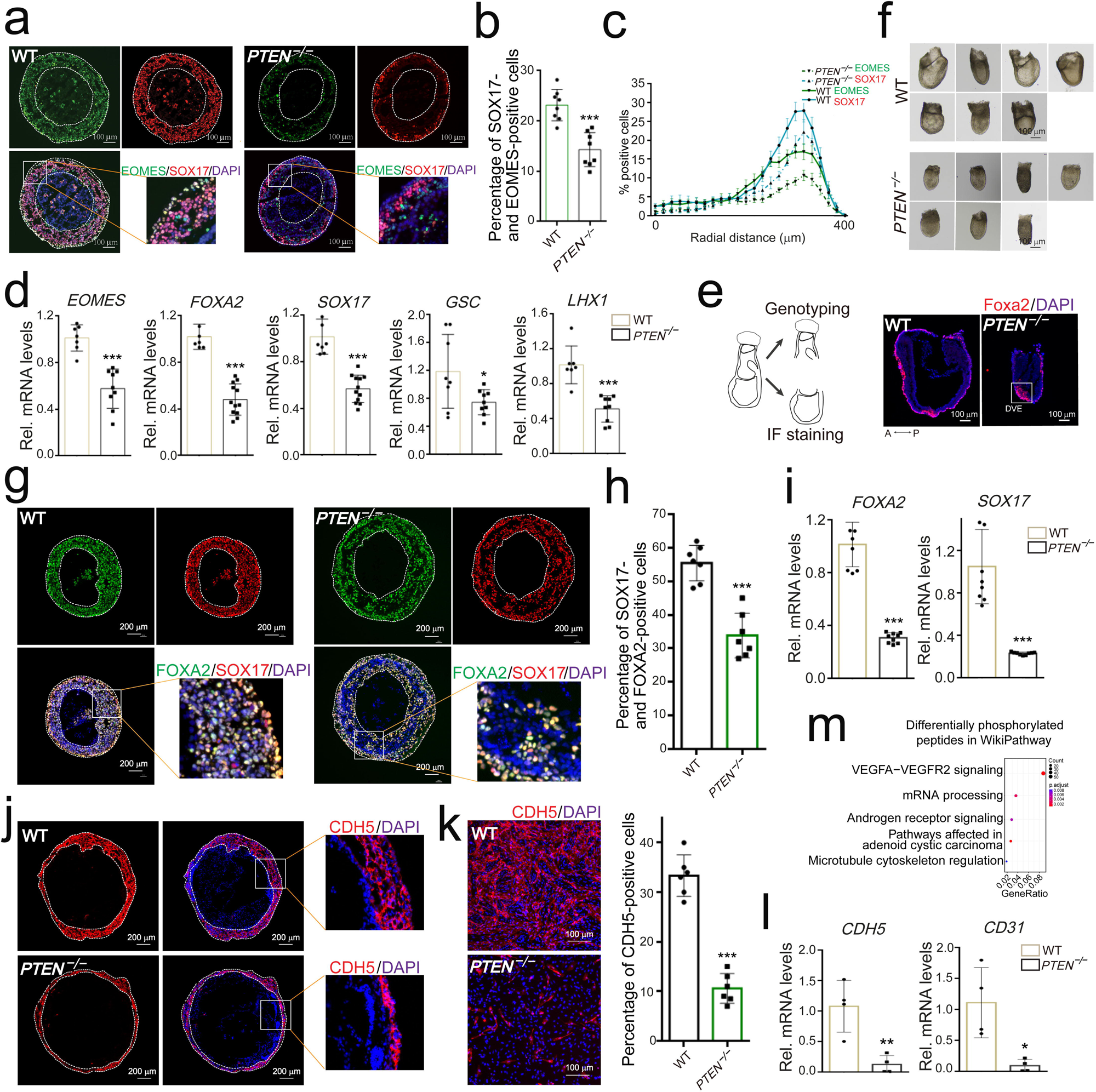
PTEN is essential for MES organoid organization and early embryonic development. **a.** *PTEN*^-/-^ MES organoids show a marked reduction in SOX17- and EOMES-positive MES cells. Scale bar, 100 µm. **b.** Proportion of MES cells co-expressing SOX17 and EOMES in WT and *PTEN*^-/-^ MES organoids. **c.** Radial profiles of the proportion of MES cells. **d.** qPCR analysis reveals significant downregulation of MES markers including EOMES, FOXA2, SOX17, GSC, and LHX1 in *PTEN*^-/-^ organoids. **e.** IF staining shows FOXA2-positive cells are predominantly localized within the distal region of the visceral endoderm in *PTEN*^-/-^ mouse embryos. Scale bar, 100 µm. **f.** Developmental progression is impaired in PTEN KO mouse embryos compared to WT at E7.5. Scale bar, 100 µm. **g.** Cryosection of a representative DE organoid demonstrates the essential role of PTEN in DE differentiation. Scale bar, 200 µm. **h.** Proportion of DE cells co-expressing SOX17 and FOXA2 in WT *andPTEN*^-/-^ DE organoids. **i.** mRNA levels of DE markers FOXA2 and SOX17 are significantly reduced in *PTEN*^-/-^ DE organoids. **j.** *PTEN^-/-^* EC organoids exhibit impaired EC ring formation. Scale bar, 200 µm. **k.** Representative images of dissociated ECs from WT *and PTEN^-/-^* organoids. Scale bar, 100 µm. **l** .*PTEN* deficiency significantly reduces mRNA expression of EC markers CDH5 and CD31. **m.** Phosphoproteomic profiling indicates enrichment of VEGFA-VEGFR2 signaling in WT versus *PTEN^-/-^* EC organoids. Statistical test, two-sided t-test; \**P*< 0.05; *\*\*P<* 0.01; ***P < 0.001.

To explore the *in vivo* relevance of these findings, we intercrossed *PTEN^+/−^* mice and collected wild-type (WT) and *PTEN^−/−^*embryos at embryonic day 7.5 (E7.5), prior to lethality. *Foxa2*, in addition to marking APS cells and DE progenitors, also orchestrates the patterning of the PS epiblast ^29^. In *PTEN^−/−^*embryos, *Foxa2*-positive cells were concentrated within the distal visceral endoderm—the precursor to the anterior visceral endoderm (AVE)—suggesting impaired cell migration (**Figure 3e**). This observation aligns with previous findings that PTEN is necessary for proper AVE migration, which determines PS positioning^30^. Moreover, *PTEN^−/−^* embryos were significantly smaller than WT counterparts, indicating delayed or defective embryonic development (**Figure 3f**).

When single cells derived from MES organoids were replated, *PTEN^−/−^*cells displayed diminished migratory capacity and failed to form converging clusters within 24 hours (Extended Data Figure 3b). These findings support a model in which PTEN promotes APS formation and MES lineage emergence during early embryogenesis.

### The generation of mesodermal and endodermal lineages is impaired by *PTEN* ablation

Gastrulation is characterized by the formation of the PS, where MES cells emerge and give rise to DE and mesoderm lineages ^16^. DE arises approximately 8 to 10 hours after the onset of gastrulation from the APS ^31^. Building on our demonstration that MES organoids can develop into DE organoids (**Figure 1h**), we investigated the role of *PTEN* in DE specification using our 3D organoid system. *PTEN* ablation significantly impaired DE generation, as evidenced by decreased *SOX17- and FOXA2*-positive cells (**Figures 3g**, 3h) and reduced expression of DE markers (**Figures 3i**).

To corroborate these findings in a 2D model, human iPSCs were directed toward DE cell differentiation, yielding *SOX17- and FOXA2*-positive cells under standard conditions. However, *PTEN* knockout markedly reduced the proportion of DE cells, as shown by IF staining (Extended Data Figures 3c, 3d), qRT-PCR (Extended Data Figure 3e), and flow cytometry (Extended Data Figure 3f). Transcriptomic analysis linked the PTEN-dependent DE defects to diabetes-related pathways (Extended Data Figure 3g). Notably, *PTEN* ablation led to diffuse nuclear localization of *SOX17* and *FOXA2 proteins* (Extended Data Figure 3c), suggesting possible nuclear fragmentation, consistent with previous studies showing that nuclear PTEN maintains chromosomal integrity ^32^. Together, these results demonstrate that PTEN is essential for MES-derived DE cell specification.

*PTEN* deletion also significantly impaired cardiac mesoderm development (Extended Data Figure 4). Both IF staining and flow cytometry revealed a decrease in the proportions of *GATA4*- and *NKX2.5*-positive cells in the *PTEN^−/−^* group (Extended Data Figures 4a, 4b). Using a defined protocol for generating human cardiac organoids (hCOs)^33^, we observed severely compromised hCO, as evidenced by diminished *NKX2.5*-positive cell populations and decreased mRNA expression levels of cardiac progenitor markers (*NKX2.5*, *MESP1*, *MESP2*) in the *PTEN^−/−^* cells (Extended Data Figures 4c–e), consistent with our previous report implicating PTEN in cardiogenesis ^34^.

ECs, which originate from mesodermal derivatives, form through vasculogenesis in both embryonic and extraembryonic tissues ^35^. When MES organoids were directed toward EC differentiation via VEGF and other factors (**Figure 1i**), *PTEN* ablation led to a marked reduction in EC ring formation (**Figure 3j**). Dissociation and replating of EC organoids further revealed significantly fewer ECs in the *PTEN^−/−^* group (**Figure 3k**). Correspondingly, mRNA levels of EC markers *CDH5* and *CD31* were significantly reduced (**Figure 3l**). Phosphoproteomic profiling indicated perturbation of the VEGFA–VEGFR2 signaling pathway (**Figure 3m**; Extended Data Figure 5a), underscoring the role of PTEN in endothelial specification from MES organoids.

### PTEN promotes MES cell generation and collective migration via its protein phosphatase activity

MES cells derived from hPSCs represent a robust *in vitro* system to model early human embryogenesis. To investigate the role of PTEN in MES generation, we employed a 2D MES differentiation protocol. *PTEN^−/−^*iPSCs showed a substantial reduction in MES cells co- expressing *EOMES* and *SOX17* (Extended Data Figures 6a-c). Rescue experiments demonstrated that PTEN overexpression (*PTEN*-OE) in the *PTEN^−/−^* iPSCs restored MES cell numbers to WT levels (Extended Data Figures 6a, 6b) and re-established expression of *EOMES*, *LHX1*, *SOX17*, *FOXA2*, and *GSC* (Extended Data Figure 6c). Bulk RNA sequencing further confirmed downregulation of MES-specific genes in *PTEN^−/−^* MES cells, without detectable expression of PPS markers (**Figure 4a**), consistent with observations made in MES organoids (**Figure 1d**; **Figure 2d**).

**Figure 4.**
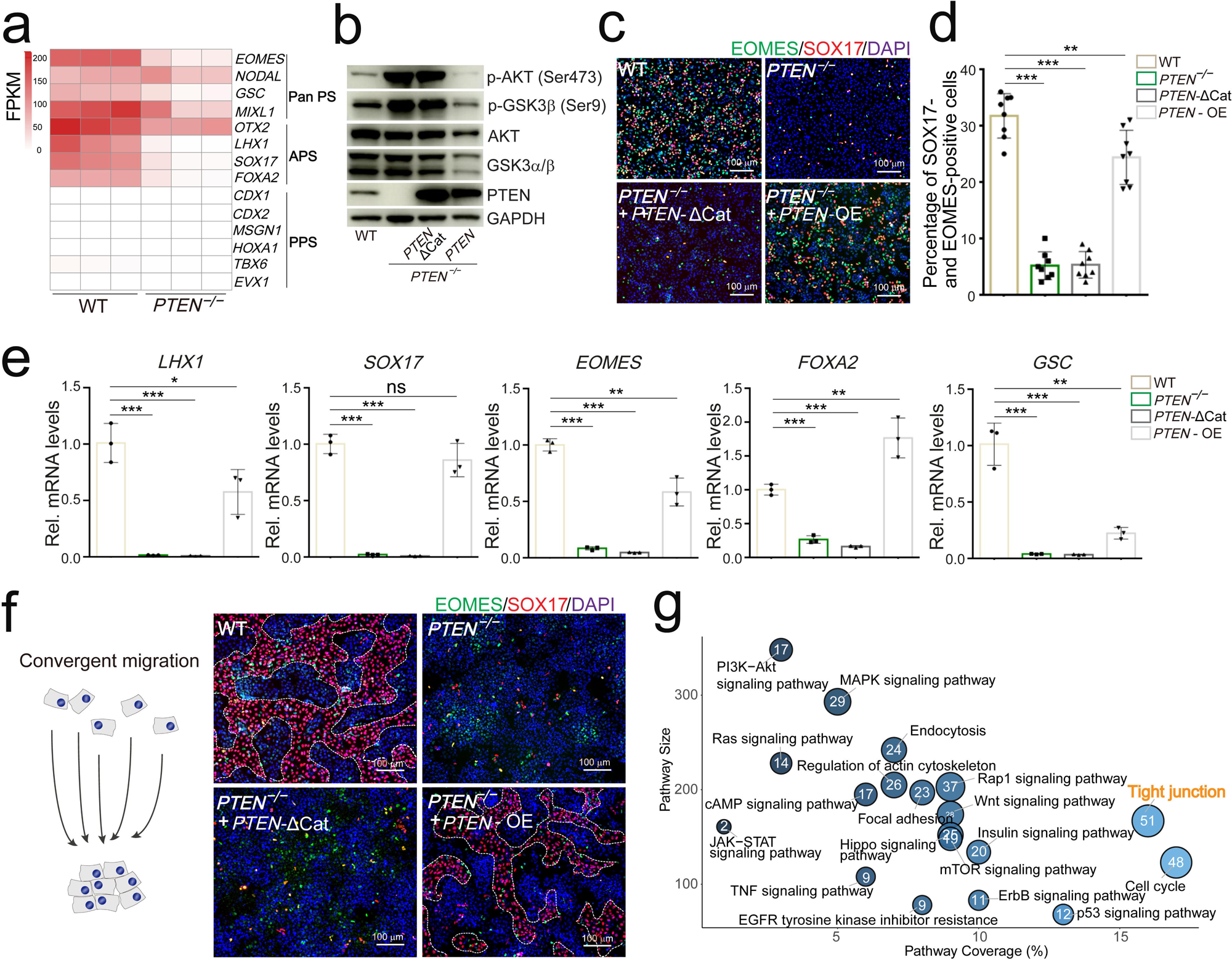
PTEN regulates MES cell generation and migration through its phosphatase activity. **a.** Heatmap showing reduced expression of pan-PS and APS markers in *PTEN^-/-^* cells, while no expression in both WT and *PTEN^-/-^* MES cells. **b.** Western blot analysis confirming loss of phosphatase activity in the catalytically inactive PTEN mutant (PTEN-ΔCat). **c, d.** PTEN-ΔCat fails to rescue MES generation *PTEN^-/-^* iPSCs. Scale bar, 100 µm. **e.** MES marker gene expression remains suppressed in *PTEN^-/-^* cells expressing PTEN-ΔCat. **f.** Convergent migration of MES cells is impaired in *PTEN^-/-^* cells and is not rescued by PTEN-ΔCat, indicating a requirement for PTEN’s enzymatic activity. Scale bar, 100 µm. **g.** Phosphoproteomic analysis reveals enrichment of kinases involved in tight junction signaling, a pathway essential for coordinated cell migration. Statistical test, two-sided t-test with Benjamini–Hochberg correction; \**P*< 0.05; \*\*\**P* < 0.001.

PTEN is best known for its lipid phosphatase activity, which antagonizes PI3K/AKT signaling ^36^. To probe whether this activity underlies its role in MES specification, we treated *PTEN^−/−^* iPSCs with the PI3K inhibitor PX-866. This treatment restored phosphorylation of AKT (Ser473) and GSK3β (Ser9) (Extended Data Figure 7a) and significantly increased the MES cell population to WT levels (Extended Data Figures 7b, 7c). MES marker expression was similarly rescued (Extended Data Figure 7d), indicating that PTEN regulates MES specification via its phosphatase activity.

To confirm this, we overexpressed a catalytically inactive *PTEN* mutant (D92A/C124A; PTEN-ΔCat) in *PTEN^−/−^* iPSCs. This mutant lacked phosphatase activity (**Figure 4b**) ^37^ and failed to restore MES generation (**Figures 4c**, 4d) or MES marker gene expression (**Figure 4e**), reinforcing that PTEN‘s phosphatase function is required for MES lineage induction.

PTEN has also been shown to promote collective cell migration in tumor models by suppressing AMPK signaling ^38^. During gastrulation, epiblast cells undergo EMT and migrate through the PS to form mesoderm and DE ^39^. Replated MES cells from *PTEN^−/−^*iPSCs exhibited markedly impaired convergence and collective migration, in contrast to WT cells (Extended Data Figure 6d). These findings underscored the critical role of PTEN in regulating this migratory behavior, indicative of the acquisition of mesenchymal characteristics. Notably, PTEN-ΔCat failed to rescue migration (**Figure 4f**), and phosphoproteomic analysis identified enrichment of migration-related tight junction pathways (**Figure 4g**).

Interestingly, *PTEN* loss also induced prominent nuclear fragmentation and promoted redistribution of *SOX17* and *EOMES* proteins from the nucleus to the cytoplasm (Extended Data Figure 6d), a phenotype similarly observed in *PTEN^−/−^* DE cells (Extended Data Figure 3c). These findings align with the established role of nuclear PTEN in preserving chromosomal integrity ^32^ and implicate nuclear-localized PTEN in the regulation of MES lineage fidelity.

### Multi-omic profiling identifies retinoic acid signalling as a PTEN-regulated pathway in mesendodermal differentiation

To investigate PTEN‘s role in MES differentiation, we induced CPC differentiation from MES cells and analyzed samples at days 3 and 6. At day 3, *PTEN^−/−^*cells exhibited significantly reduced MES marker expression compared to WT, with both groups showing diminished MES marker levels by day 6 (**Figure 5a**). However, CPC marker expression was markedly lower in *PTEN^−/−^* cells at day 6, with no detectable MES gene expression (**Figure 5b**). Transcriptional profiling across differentiation stages revealed distinct gene expression trajectories between WT and *PTEN^−/−^* groups. PCA demonstrated clear clustering by genotype and timepoint (**Figure 5c**), and pathway enrichment analysis identified significant downregulation of signaling pathways involved in organogenesis and anterior-posterior (A-P) patterning in *PTEN^−/−^* cells at day 3 (**Figure 5d**), consistent with PTEN‘s established role in early lineage commitment from iPSCs.

**Figure 5.**
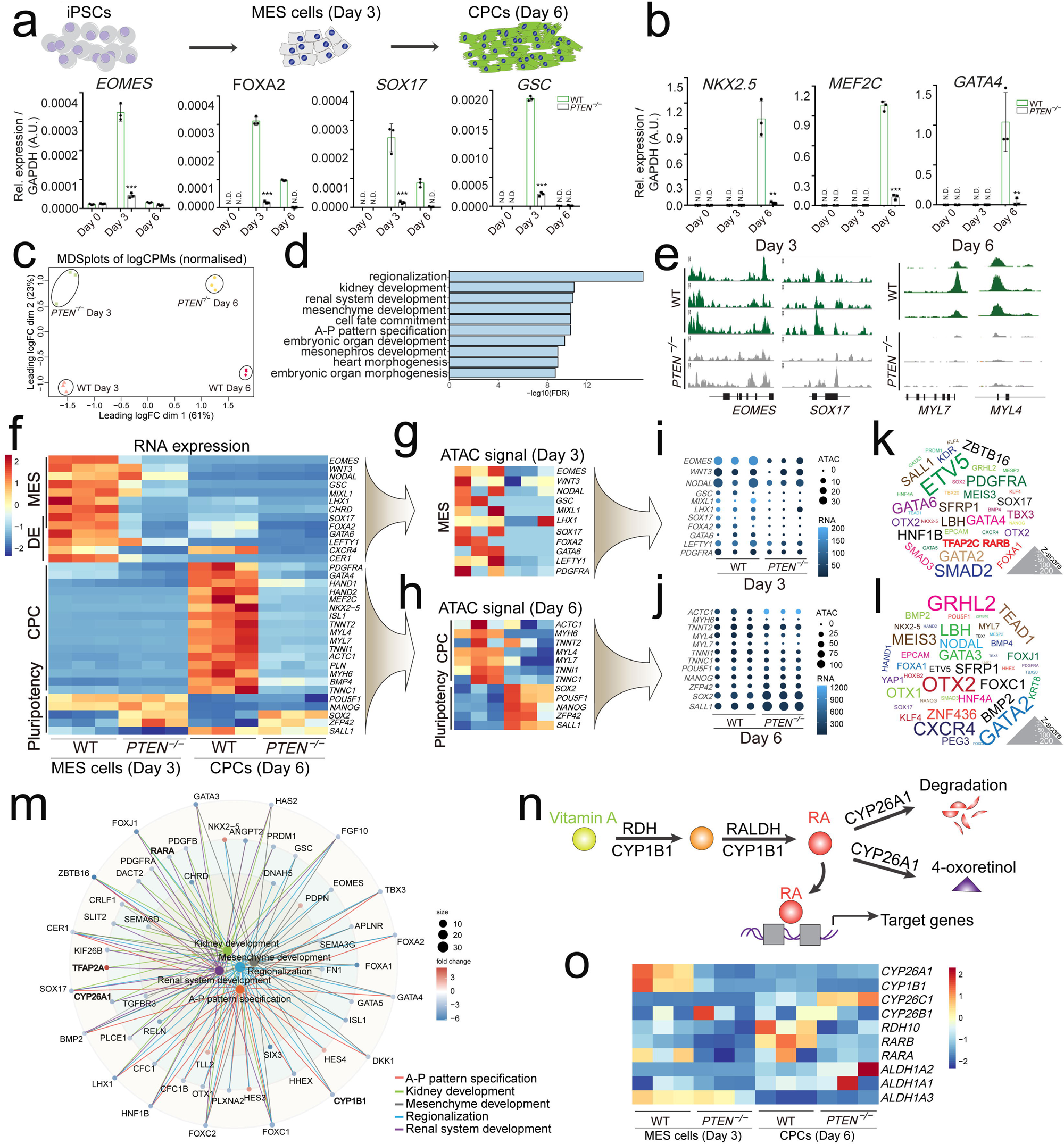
Multi-omic analyses identify RA signaling regulators in MES cell generation. **a, b.** mRNA expression of MES and CPC markers at days 3 and 6, respectively, showing that *PTEN* ablation disrupts their temporal induction. **c.** PCA of transcriptomic data separates WT and *PTEN^-/-^* samples at both differentiation stages. **d.** GO enrichment analysis of RNA-seq data highlights pathways involved in mesodermal and endodermal lineage specification. **e.** ATAC-seq reveals that PTEN loss reduces chromatin accessibility at MES marker loci (SOX17, EOMES) at day 3 and CPC marker loci (MYL7, MYL4) at day 6. **f.** Heatmap showing expression of APS, DE, CPC, and pluripotency markers in WT versus *PTEN^-/-^* MES and CPC populations. **g-1.** Integrated RNA-seq and ATAC-seq analysis identifies enrichment of RA signaling components and transcription factors involved in gastrulation and cardiac differentiation in WT cells. **m.** Pathway network analysis revealing significant enrichment of key RA signaling regulators in WT compared *toPTEN^-/-^ MES* cells. **n, o.** Heatmaps indicate that CYP26A1, a key RA catabolic enzyme, is significantly downregulated in *PTEN^-/-^* MES cells. Statistical test, two-sided t-test; *\*P* < 0.05; *\*\*P* < 0.01; *\*\*\*p<* 0.001.

Chromatin accessibility profiling using ATAC-seq revealed that *PTEN* deletion markedly reduced accessibility at the loci of MES-specific genes, such as *SOX17* and *EOMES*, at day 3, and CPC markers, including *MYL7* and *MYL4,* at day 6 (**Figure 5e**; Extended Data Figure 6e). Complementary transcriptomic analysis showed significant downregulation of MES and DE markers in *PTEN^−/−^* cells at day 3, followed by a decline in cardiac lineage gene expression and a concurrent upregulation of pluripotency-associated genes at day 6 (**Figure 5f**). These results are consistent with previous findings implicating PTEN in both cardiac differentiation and pluripotency regulation ^34, 40^. The ATAC-seq data further supported these transcriptomic trends, demonstrating reduced chromatin accessibility at MES and cardiac loci in *PTEN^−/−^* cells, alongside increased accessibility at pluripotency gene regions (**Figures 5g**, 5h). A bubble plot analysis reinforced these findings, showing that *PTEN* ablation significantly diminished accessibility at MES gene loci on day 3 and CPC gene loci on day 6, while enhancing accessibility at loci associated with pluripotency (**Figures 5i**, 5j).

Motif enrichment analysis revealed that WT MES cells preferentially enriched for TFs critical to gastrulation, including *ETV5*, *SMAD2*, *GATA2*, *SALL1*, *PDGFRA*, *SFRP1*, *SOX17*, *HNF1B*, *GATA4*, and *GATA6* (**Figure 5k**). Additionally, cardiac-specific TFs such as *OTX2*, *HAND1*, *NKX2.5*, *OTX1*, *LBH*, *TEAD1*, and *MEF2C* were significantly enriched in WT cells relative to *PTEN^−/−^*CPCs at day 6 (**Figure 5l**). These results collectively indicate that PTEN is essential for maintaining chromatin accessibility at MES and CPC gene loci, aligning with the transcriptomic data (**Figures 5f**).

In addition to developmental regulators, pathway network analysis identified key components of the retinoic acid (RA) signaling cascade—*RARA*, *CYP26A1*, *CYP1B1*, and *TFAP2A*—as significantly enriched in WT compared to *PTEN^−/−^*MES cells (**Figure 5m**). Heatmap visualization of RA pathway regulators revealed robust expression of CYP26A1 in WT MES cells, whereas no significant difference was observed between WT and *PTEN^−/−^* CPCs (**Figures 5n**, 5o). CYP26A1 is essential for spatially restricting RA signaling during embryogenesis^41^. Together, these results suggest that RA signaling is a critical downstream effector of PTEN in regulating MES cell generation.

### CYP26A1 functions downstream of PTEN in MES specification

A volcano plot analysis of transcriptomic data revealed that *CYP26A1*, along with multiple MES marker genes, was significantly downregulated in *PTEN^−/−^*MES cells (**Figure 6a**), a result corroborated by ATAC-seq, which showed decreased chromatin accessibility at these loci (**Figure 6b**). Western blotting confirmed a marked reduction in *CYP26A1* protein levels following PTEN loss (**Figure 6c**). In comparison, *PTEN*-OE restored *CYP26A1* expression at both the mRNA and protein levels (**Figures 6c**, 6d), establishing *CYP26A1* as a downstream effector of PTEN. Importantly, overexpression of *CYP26A1* (*CYP26A1*-OE) in *PTEN^−/−^* iPSCs fully rescued the population of MES cells co-expressing *SOX17* and *EOMES* (**Figures 6e**, 6f), and reinstated the expression of MES markers suppressed by *PTEN* deletion (**Figure 6g**). Moreover, impaired MES cell migration induced by *PTEN* loss was completely rescued by *CYP26A1*-OE, restoring migration to WT levels (**Figure 6h**). These findings support previous studies showing that PTEN is essential for epiblast migration during mouse gastrulation^21^. Conversely, *CYP26A1* knockdown via shRNA significantly impaired MES cell generation and downregulated MES markers (Extended Data Figures 8a–c). Additionally, we generated *CYP26A1^−/−^* iPSCs and observed that the ablation of *CYP26A1* significantly inhibited MES formation by reducing the proportion of MES cells as well as the mRNA levels of MES marker genes (Extended Data Figures 8d–f ). As anticipated, MES organoids derived from *CYP26A1^−/−^* iPSCs also demonstrated a substantial decrease in *SOX17*- and *EOMES*-positive cells (**Figure 6i**, 6j), which is fully consistent with the findings from *PTEN^−/−^* iPSCs (**Figure 3a**, 3b). A kinome dendrogram from phosphoproteomic analysis revealed widespread alterations in kinase activity following *PTEN* deletion, including WNT signalling (**Figure 6k**). Notably, phosphoproteomic profiling demonstrated significant suppression of β-catenin signaling in *PTEN^−/−^* cells (**Figure 6l**), consistent with the established role of stabilized β-catenin in inducing *CYP26A1* expression ^42^. Additionally, WNT/β-catenin signaling was further validated by Western blot and transcriptome data analyses (**Figure 6m**, 6n).

**Figure 6.**
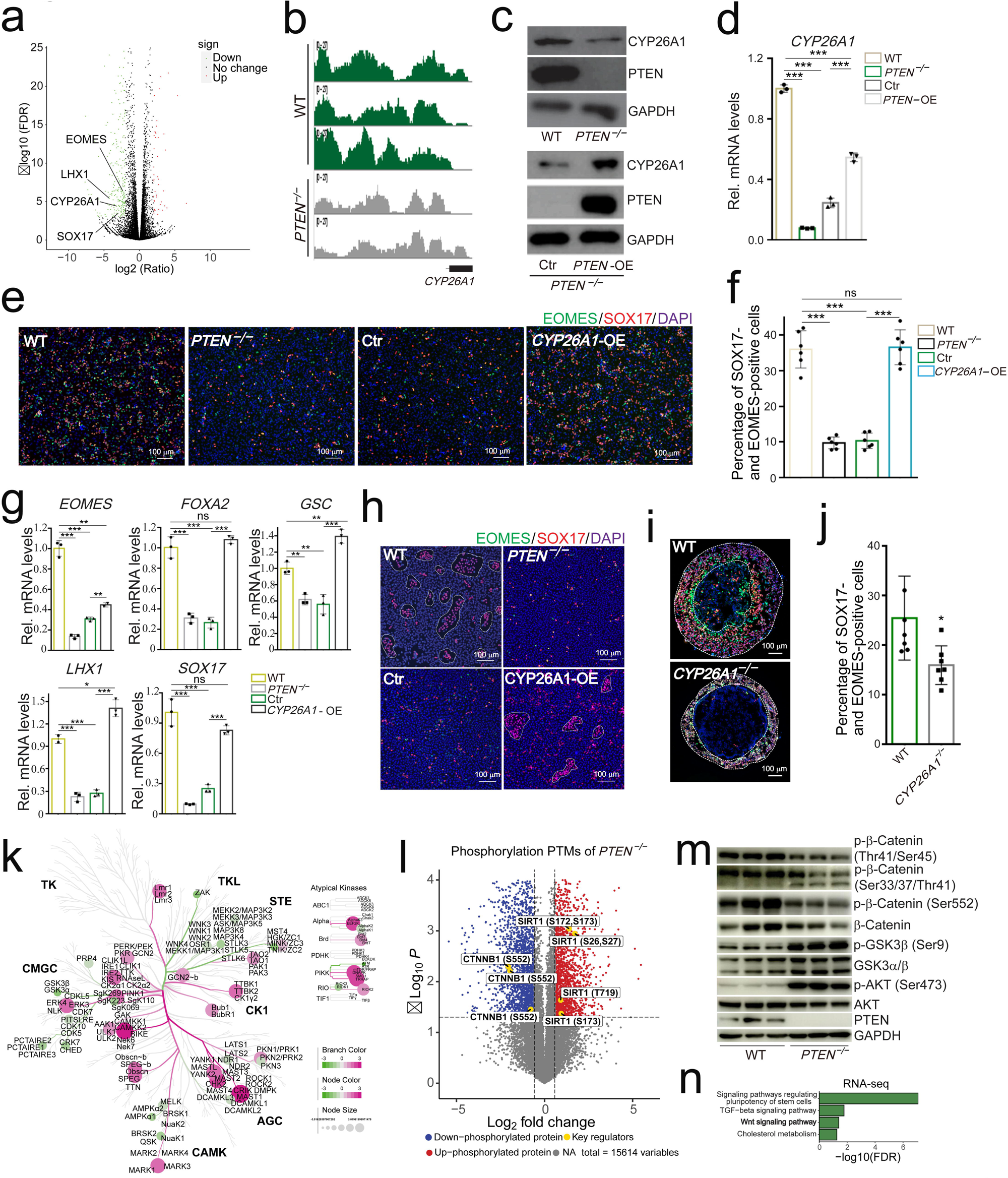
PTEN regulates CYP26A1 expression during MES lineage specification. **a.** Volcano plot showing differentially expressed genes between WT and PTEN*^-/-^* MES cells; CYP26A1 and MES markers are notably downregulated in thePTEN*^-/-^* group. **b.** ATAC-seq reveals reduced chromatin accessibility at the CYP26A1 locus in cells. **c, d.** PTEN-OE restores CYP26A1 expression at both transcript and protein levels. **e, f.** CYP26A1-OE in *PTEN^-/-^* iPSCs rescues MES cell generation, as evidenced by restored co-expression of SOX17 and EOMES. Scale bar, 100 µm. **g.** The expression levels of MES markers are rescued following CYP26A1-OE in PTEN^_/_^ cells. **h.** CYP26A1-OE fully rescues MES cell migration impaired by PTEN loss. Scale bar, 100 µm. **i.** *CYP26A1^-/-^* MES organoids show a marked reduction in SOX17- and EOMES-positive MES cells. Scale bar, 100 µm. **j.** Proportion of MES cells co-expressing SOX17 and EOMES in WT and *CYP26A1 ^-/-^* MES organoids. **k.** Kinome dendrogram derived from human phosphorylation proteomics data illustrating kinase signaling pathways enriched in WT and *PTEN^-/-^* MES cells. **l.** Phosphoproteomic profiling reveals reduced phosphorylation of β-catenin (Ser552) and increased phosphorylation of SIRT1 in *PTEN^-/-^* cells. **m** Western blot analysis shows increased AKT and GSK3β phosphorylation and decreased β-catenin phosphorylation in *PTEN^-/-^* cells. **n.** WNT signaling pathways are significantly enriched in the WT MES transcriptome. Statistical test, two-sided t-test with Benjamini–Hochberg correction; *\*P<* 0.05; \*\*\**P*< 0.001.

Furthermore, treatment of *PTEN^−/−^* cells with the PI3K inhibitor PX-866 elevated total and phosphorylated β-catenin (Ser552) and restored *CYP26A1* expression (Extended Data Figure 9a). Both PX-866 and *PTEN*-OE, but not catalytically inactive PTEN (PTEN-ΔCat), rescued MES marker expression (Extended Data Figures 9b, 9c). Heatmap analysis and kinome profiling further confirmed the upregulation of WNT signaling regulators (Extended Data Figures 9d, 9e). Collectively, these data suggest that PTEN drives MES specification via *CYP26A1*, acting through phosphatase-dependent mechanisms that converge on RA and WNT signalling pathways.

### PTEN facilitates MES specification by regulating RA homeostasis

RA is synthesized in distinct embryonic regions and regulates transcription by binding nuclear RA receptors (RARs), which in turn interact with RA response elements (RAREs) near target genes. However, excessive RA is known to be teratogenic ^43^. CYP26A1, an RA- inducible cytochrome P450 enzyme, degrades RA into polar metabolites such as 4-hydroxy- RA and 4-oxoretinol (4-oxo-RA) ^44^. CYP26A1 plays a critical role in maintaining RA homeostasis by depleting excessive RA within specific embryonic regions, thereby shaping the A-P axis.

To examine whether RA participates in PTEN‘s effect on MES generation, we performed lipid metabolite profiling focused on gastrulation-associated metabolites. Notably, 4-oxo-RA levels were significantly decreased in *PTEN^−/−^* MES cells (**Figure 7a**), a finding confirmed by heatmap analysis showing diminished 4-oxo-RA upon *PTEN* deletion (**Figure 7b**). These results suggest that loss of *PTEN* reduces *CYP26A1* expression, impairing RA catabolism and resulting in RA accumulation in *PTEN^−/−^* MES cells.

**Figure 7.**
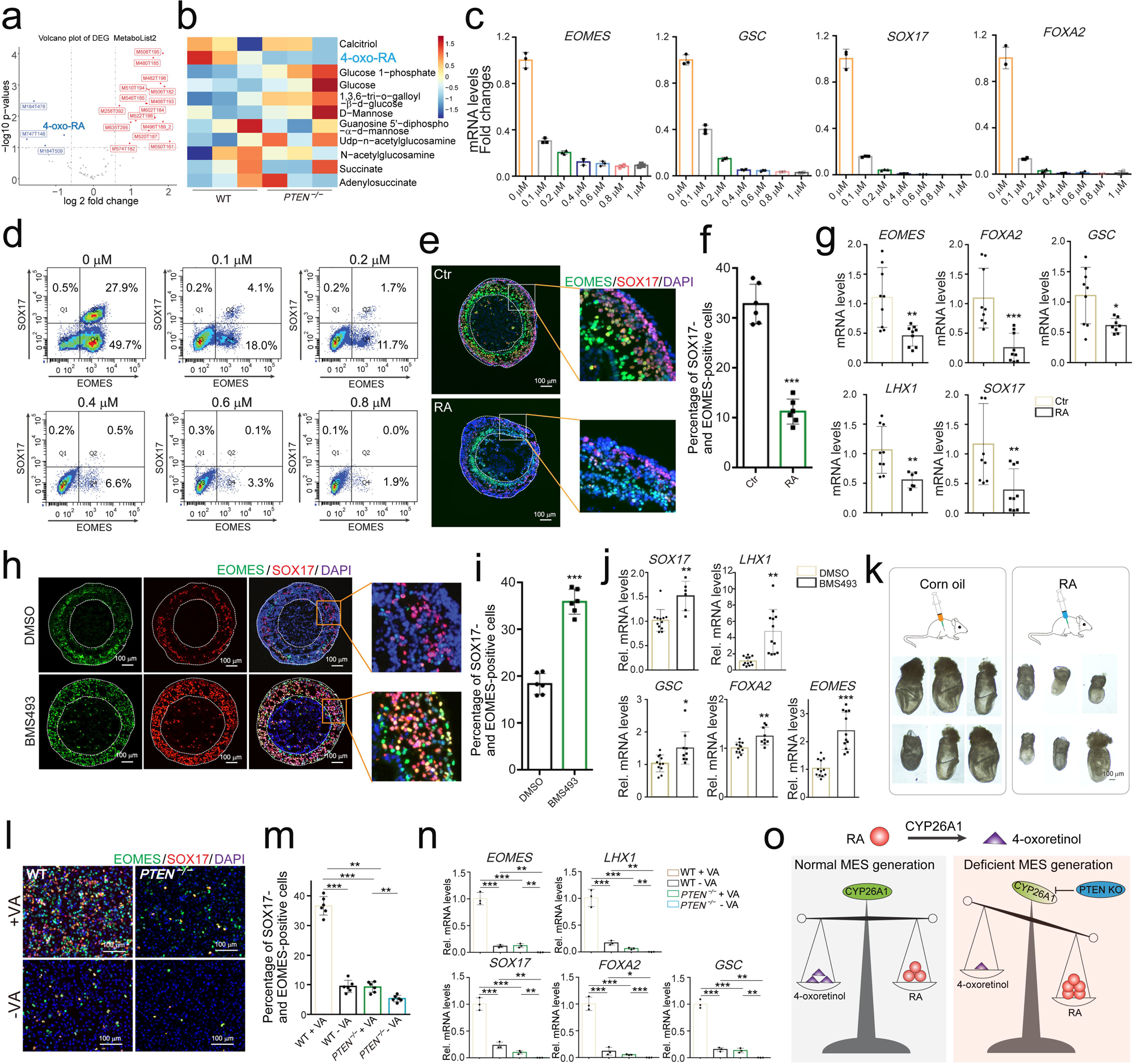
Both excessive and insufficient RA impairs MES differentiation. **a.** Dot plot of metabolomics data shows significant reduction of 4-oxo-RA among all the in *PTEN^-/-^* MES cells. **b.** Heatmap profiling of gastrulation-related metabolites confirms marked depletion of 4-oxo-RA following *PTEN* ablation. **c.** qPCR analysis reveals dose-dependent suppression of MES marker genes in WT cells upon exogenous RA treatment. **d.** Flow cytometry shows reduced MES cell proportions following RA exposure. **e, f.** RA treatment reduced SOX17- and EOMES-positive cells in MES organoids. Scale bar, 100 µm. **g.** qPCR confirms downregulation of MES markers in RA-treated MES organoids. **h, i.** MES organoid assays confirm that RA antagonist BMS493 enhances MES formation. Scale bar, 100 µm. **j.** qPCR data show robust upregulation of MES markers following RA inhibition. **k.** Morphological comparison of control and RA-treated embryos illustrates developmental delays with excess RA. Scale bar, 100 µm. **1, m.** IF staining of MES cell formation under vitamin A-supplemented versus vitamin A-depleted conditions. Scale bar, 100 µm. **n.** qPCR analysis demonstrates that both vitamin A and RA are essential, but tightly regulated, for proper MES cell differentiation. **o.** Schematic model proposing that CYP26A1-mediated RA degradation is critical for maintaining RA homeostasis during MES specification. Statistical test, two-sided t-test; *\*P <* 0.05; *\*\*p<* 0.01; ***P<0.001.

We hypothesized that elevated RA levels suppress MES differentiation in *PTEN*-deficient cells. To test this, WT iPSCs were treated with 1 µM RA, resulting in a significant reduction in the expression of MES markers, closely recapitulating the expression profile observed in *PTEN^−/−^*MES cells (Extended Data Figure 10a). RA exposure led to a dose-dependent inhibition of MES differentiation (Extended Data Figure 10b), with progressive downregulation of MES markers including *EOMES*, *FOXA2*, *SOX17*, and *GSC,* mirroring the expression profile observed in *PTEN^−/−^* MES cells (**Figure 7c**). Flow cytometry confirmed reduced MES cell proportions following RA treatment (**Figure 7d**). Similarly, in MES organoids, RA treatment decreased MES cell populations and marker expression (**Figures 7e–g**; Extended Data Figure 10c).

To confirm the inhibitory role of RA in MES specification, we treated *PTEN^−/−^* iPSCs with the pan-RAR antagonist BMS493. BMS493 significantly increased MES cell proportions (Extended Data Figure 10d) and partially restored MES gene expression (Extended Data Figure 10e). In *PTEN^−/−^* MES organoids, BMS493 treatment similarly enhanced the number of SOX17- and EOMES-positive cells as well as the expression of MES markers (**Figures 7h–j**; Extended Data Figure 10f). *In vivo*, pregnant mice treated with RA at E4 yielded E7.5 embryos that were smaller (**Figures 7k**), phenocopying *PTEN^−/−^* embryonic phenotypes (**Figures 3f**).

RA signaling is essential for the development of multiple organ systems, including the axial skeleton, spinal cord, heart, and reproductive tract ^45^. Recent studies also show that vitamin A, the RA precursor, restricts lineage plasticity and guides stem cell fate decisions ^46^, and that RA induces posterior embryonic structures in human gastruloids ^47^. Consistently, withdrawal of vitamin A from the MES differentiation medium markedly suppressed MES generation in both WT and *PTEN^−/−^*cells, as evidence by reduced numbers of SOX17- and EOME*S*- positive cells and decreased expression of MES markers (**Figures 7l–n**), underscoring the essential role of RA in MES lineage specification. Collectively, these data indicate that both RA excess and deficiency impair MES differentiation, and that *CYP26A1* maintains RA homeostasis critical for proper embryonic patterning (**Figure 7o**).

## Discussion

Defects arising from aberrant gastrulation are frequently incompatible with life and are a major cause of spontaneous miscarriage. The incidence of spontaneous abortion ranges from 10% to 15%, with many cases linked to genetic anomalies that may contribute to congenital growth and developmental disorders ^48^. Specification of MES at the PS is a transient yet critical developmental milestone. Due to the inaccessibility and complexity of early human embryos, our understanding of native human gastrulation remains limited, highlighting the need for sophisticated *in vitro* models. Here, we establish an organoid system that faithfully recapitulates human MES lineage specification (**Figure 1a-f**). These organoids can be further differentiated into EC and DE organoids (**Figure 1g-i**), which may serve as precursors for vascular, hepatic, or pancreatic tissues. Collectively, this system provides a valuable platform for dissecting early lineage specification, modeling congenital disorders, and facilitating the discovery of drugs targeting developmental abnormalities.

*PTEN* loss-of-function mutations are known to cause embryonic lethality, and aberrant *PTEN* expression has been associated with spontaneous abortion in clinical settings ^22, 23, 24^. RA signaling plays a crucial role in embryogenesis, including cell-cell communication and organ formation ^20^, but dysregulated RA levels—whether excessive or deficient—can impair early development, especially blastocyst formation and implantation^49^. Although RA is essential for early patterning, its role in PS formation, particularly in the specification of APS- derived lineages, remains unclear ^20^. Our study revealed that *PTEN^−/−^* MES cells exhibit significantly reduced levels of 4-oxo-RA (**Figure 7a**, 7b), a CYP26A1-catalyzed RA metabolite, suggesting elevated intracellular RA. These elevated RA levels suppress MES lineage specification (**Figures 7c–g**; Extended Data Figure 10a–c), while RA deficiency similarly disrupts MES differentiation (**Figures 7l–o**), aligning with clinical observations of RA‘s dose-sensitive teratogenicity ^50, 51^.

Building on these findings, we propose a model in which *PTEN* ablation suppresses MES cell generation by dysregulating CYP26A1-mediated RA homeostasis. Vitamin A or RA deficiency during pregnancy can cause fetal defects such as night blindness, skeletal malformations, and epithelial dysplasia, while excessive intake is associated with severe central nervous system abnormalities ^51^. Since *PTEN*-null embryos fail to survive beyond E7.5 ^21^, we propose that *PTEN* loss leads to RA imbalance, contributing to the observed developmental arrest. Our data suggest that PTEN maintains appropriate RA levels by regulating *CYP26A1*, thereby enabling proper MES lineage specification (**Figure 7o**).

RA signaling has also been shown to promote self-renewal of mouse embryonic stem cells via activation of the PI3K/AKT pathway ^52^. However, our results showed that PTEN may regulate CYP26A1 expression and RA signaling. Our data indicate that *PTEN* loss suppresses β-catenin protein levels and its phosphorylation, which may disrupt CYP26A1 transcription (**Figures 6l**, 6n). Notably, the phosphorylation sites affected (Thr41/Ser45/Ser33/Ser37) are linked to β-catenin degradation, suggesting a complex mechanism by which PTEN stabilizes β-catenin. *PTEN^−/−^* MES cells also exhibited elevated phosphorylation of SIRT1, a NAD- dependent deacetylase, at S26, S27, S172, and S173—modifications associated with increased deacetylase activity (**Figure 6l**). Given SIRT1‘s role in regulating stem cell fate through β-catenin deacetylation ^53, 54^, these findings suggest that enhanced SIRT1 activity in *PTEN^−/−^* cells may destabilize β-catenin via increased deacetylation (Extended Data Figures 9f). Supporting this, *PTEN* loss resulted in decreased β-catenin protein levels and reduced phosphorylation at Ser552 as well as an increased phosphorylation level of SIRT1 at S27, effects likely attributable to SIRT1-mediated destabilization (**Figure 6l**). We hypothesize that SIRT1 may destabilize β-catenin through increased deacetylation in *PTEN*-deficient cells, while further experimental validation is needed to confirm this hypothesis (Extended Data Figures 9f).

In addition to its role in RA homeostasis, PTEN promotes MES cell migration in a phosphatase-dependent manner (**Figure 4f**), in agreement with prior studies ^55^. We also observed nuclear fragmentation upon *PTEN* deletion during MES migration (Extended Data Figure 6f), suggesting that nuclear structural integrity may be essential for cell motility.

Moreover, AKT1 signaling is known to promote the transition from MES to anterior DE ^56^, and PTEN has been implicated in vasculogenesis during early chick development ^57^. As both DE and EC organoids are derived from MES organoids, the observed impairment in their formation following *PTEN* ablation reinforces the essential role of *PTEN* in lineage progression toward endodermal and endothelial fates, consistent with these prior reports ^56, 57^. This model thus serves as a robust platform for studying lineage commitment during early human development.

Deciphering the molecular mechanisms by which PTEN orchestrates MES specification and APS formation enhances our understanding of the pathogenesis underlying certain congenital malformations and spontaneous miscarriages. This work identifies CYP26A1-mediated RA signaling as a central node in *PTEN*‘s function, providing novel insights into the regulatory networks potentially driving human gastrulation and PS formation. These findings also offer a molecular basis for investigating congenital abortion and open avenues for therapeutic strategies aimed at preventing early pregnancy loss.

## Methods

### Cell culture, MES cell generation, MES organoid formation

Human iPSCs (Allen Institute for Cell Science, AG28869) were cultured using mTeSR1 medium (STEMCELL Technologies, 85850). For 2D MES cell generation, hPSCs were seeded at a density of 250,000 cells/well in 12-well plate in mTeSR1+ROCKi (Y27631, Selleckchem, S1049). After 48 hours of culture, MES differentiation was initiated by replacing the medium with RPMI/1640 medium (Life Technologies, 11875085) supplemented with 2% B-27 without insulin (Life Technologies, A1895601) and 6 µM CHIR990021 (Selleckchem, S1263). Following 48 hours of induction, the medium was replaced with RPMI/1640 medium containing 2% B-27 supplement without insulin for 24 hours. Cells were then harvested for Q-PCR, Western blot, and immunofluorescence (IF) staining.

For MES organoid formation, hPSCs were seeded at 5,000 cells/well in a U-shaped ultralow- attachment 96-well plate (Corning, 3788) in mTeSR1+ROCKi. The plates were centrifuged at 300 ×g and incubated to facilitate aggregate formation. After 24 hours, the cell aggregates were embedded in Matrigel droplets (Corning, 354234). The plates were incubated for 1 hour to allow Matrigel solidification, after which mTeSR1 medium was added. Differentiation was initiated on d 0 by replacing the medium with RPMI/1640 medium containing 2% B-27 supplement without insulin and 7.5 µM CHIR990021. After 24 hours, the medium was replaced with RPMI/1640 medium containing 2% B-27 without insulin. Following an additional 48 hours, MES organoids were collected for downstream analyses.

### DE cell generation from hiPSCs

hPSCs were seeded at 250,000 cells/well in 12-well plates in mTeSR1+ROCKi. After 48 hours of culture, MES differentiation was initiated by replacing the medium with RPMI/1640 medium supplemented with 2% B-27 without insulin, 3 µM CHIR990021, and 100 ng/ml Activin A (R & D Systems, 338-AC-050). After 24 hours, the medium was replaced with RPMI/1640 medium containing 2% B-27 supplement and 100 ng/ml Activin A for an additional 2 days. Subsequently, DE cells were harvested for Q-PCR, flow cytometric analysis, and IF staining.

### DE and EC organoid formation

After collection of the MES organoids, differentiation of DE and EC organoids as initiated. For DE organoid generation, the culture medium was replaced with RPMI/1640 medium supplemented with 2% B-27 without insulin and 100 ng/ml Activin A for an additional 3 days. For EC organoid generation, MES organoids were cultured in RPMI/1640 medium containing 2% B-27 without insulin, 100 ng/ml VEGF (PeproTech, 100-20), and 2 µM forskolin (Selleckchem, S2449) for 48 hours. Subsequently, the medium was exchanged to RPMI/1640 medium containing 2% B-27 supplement, 100 ng/ml VEGF, 100 ng/ml FGF-2 (PeproTech, AF-100-18B), and 15% fetal bovine serum for an additional 5 days.

### Laboratory animals

All mice were maintained in a specific pathogen-free environment, and all animal experiments were conducted in accordance with the guidelines and regulations of Shandong University. The experimental protocol was approved by the Animal Care and Research Committee of Shandong University.

### Genome edited iPSC generation

PTEN and CYP26A1 deletion in human iPSC were generated as previously detailed ^34, 58^. Guide RNA sequences were incorporated into the pSpCas9 (BB)-2A-GFP vector (Addgene, 48138), which encodes both Cas9 nuclease and green fluorescent protein (GFP). iPSCs were cultured in 6-well plates and transfected with the Cas9-sgRNA constructs using Lipofectamine 3000 (Invitrogen). Two days post-transfection, GFP-positive cells were isolated by fluorescence-activated cell sorting (FACS). The enriched, gene-edited cell populations were subsequently cultured in 48-well plates for 4-5 days, followed by dilution and plating into 96-well plates to enable single-colony formation. Individual colonies were expanded and screened for *PTEN^−/−^* and *CYP26A1^−/−^* genotypes via DNA sequencing. Primer sequences used for *CYP26A1* deletion are provided in **Supplementary Table** 1.

### Plasmid Construction

Human *CYP26A1* genes were amplified by RT-PCR from total RNA isolated from human iPSCs and subsequently cloned into the pCDH-EF1-MCS-BGH-PGKcopGFP-T2A-Puro Vector (System Biosciences, CD550A-1). The primer sequences are provided in **Supplementary Table** 1. The *PTEN*-OE and *PTEN*-ΔCat-OE system were previously described ^34^. To achieve knockdown of *CYP26A1* expression, shRNA constructs were generated and cloned into pKLO.1 vector. The shRNA oligonucleotide sequences are listed in **Supplementary Table** 1.

### Cryosectioning and staining

Organoids were harvested and subsequently fixed in 4% PFA. Following fixation, samples were permeabilized with 0.3% Triton X-100 for 20 minutes. Following washing with PBS, organoids were stored at 4°C until further processing. For cryo-embedding, organoids were transferred into tissue embedding molds containing OCT and rapidly frozen at low temperatures. Tissue sections of 10 μm thickness were then prepared using either the Epredia Cryostar NX70 Cryostat or the Leica CM1950 Cryostat.

### Immunofluorescent staining

The cells were fixed in 4% PFA for 20 minutes at room temperature, followed by three washes with PBS for 5 minutes each. Subsequently, the samples were incubated with permeabilization buffer and washed three additional times with PBS. To minimize non- specific binding, the samples were blocked with a blocking buffer for 20 minutes at room temperature. Thereafter, the cells were incubated overnight at 4 °C with primary antibodies diluted in staining buffer. The following day, samples were incubated with secondary antibodies for 1 hour at room temperature in the dark. Finally, fluorescence imaging was performed using a fluorescence microscope. Primary antibodies were applied at the following dillutions: anti-SOX17 (R & D Systems, AF1924, 1:1000), anti-EOMES (Cell Signaling Technology, 81493, 1:1000), anti-FoxA2 (Cell Signaling Technology, 8186, 1:1000), and anti-CDH5 (Cell Signaling Technology, 2500, 1:1000).

### Flow cytometry

Both 2D cultured cells and 3D organoids were collected into tubes and digested into single- cells suspensions using Accutase (Sigma, A6964) for 10 minutes at 37°C. The digested cells were fixed with 4% PFA for 10-15 minutes. Following fixation, cells were washed with 500 µl of FACS buffer and centrifuged at 1500 ×rpm for 5 minutes at 4°C. The supernatant was carefully aspirated. Cells were then permeabilized by incubation with 200 µl of 0.3% Triton- X100 buffer at room temperature for 15 minutes. After an additional wash with 500 µl FACS buffer and centrifugation at 2000 rpm for 5 minutes at 4°C, the supernatant was discarded gently. Subsequently, cells were incubated on ice with primary antibodies (Anti-SOX17 antibody, anti-EOMES antibody; Anti-SOX17 antibody, anti-FoxA2 antibody) for 45-60 minutes. Following primary antibody staining, cells were washed with 500 µl FACS buffer and centrifuged under the same conditions. Cells were then resuspended in 100 µl of secondary antibodies, mixed thoroughly, and incubated at room temperature for 20-30 minutes in the dark. After a final wash and centrifugation, cells were resuspended in FACS buffer and filtered with Falcon 5 ml round bottom polystyrene test tube to prepare for flow cytometry analysis.

### Western blot

Protein samples were prepared by lysing cells in RIPA buffer supplemented with proteinase inhibitor for 20 minutes on ice. Lysates were then centrifuged at 12000 rpm for 5 minutes at 4°C, and the supernatant was collected. The protein extracts were mixed with 5×SDS sample buffer and denatured by heating at 98°C for 10 minutes. Samples were loaded onto SDS- PAGE gels and electrophoresed initially at 80 V, followed by 120 V until the dye front reached the bottom of the gel.

Following electrophoresis, polyvinylidene difluoride (PVDF) membranes were equilibrated by soaking in transfer buffer for 1–2 minutes and briefly rinsed. The gel and membrane were assembled in a transfer apparatus, and proteins were transferred at 110 V for 75 minutes. Subsequently, membranes were blocked in blocking buffer for 1 hour at room temperature.

Primary antibodies diluted in blocking buffer were incubated with the membranes overnight at 4°C. After incubation, membranes were washed three times with TBST for 10 minutes each. Membranes were then incubated with secondary antibodies diluted in blocking buffer for 1 hour at room temperature, followed by three additional washes with TBST for 10 minutes each. The membranes were then ready for imaging and further analysis.

### Real-time PCR Analysis

Total RNA was extracted from collected cells using TRIzol reagent (Invitrogen) according to the manufacturer‘s instructions. Approximately 400 ng of RNA was reverse-transcribed into complementary DNA (cDNA). Quantitative PCR (qPCR) was performed using SYBR Green chemistry, and relative mRNA expression levels were calculated using the ΔΔCt method. Gapdh was selected as the endogenous reference gene for normalization. All qPCR assays were conducted with technical replicates to ensure reproducibility. Primer sequences are provided in **Supplementary Table** 1.

### Image processing and data quantification

IF staining images were analyzed using custom scripts in ImageJ. Cell counting was performed utilizing the StarDist plugin for ImageJ. Three-dimensional reconstruction and volumetric analysis of MES and EC organoids were conducted using the 3D ImageJ Suite.

### ScRNA-seq for MES organoids

MES organoids were collected on d3 and individually dissociated into single cells using Accutase. The dissociated cells were washed with PBS and centrifuged for 5 minutes; the cells were resuspended ensuring that there were above 100 × 10^3^ cells for each sample. Library preparation for scRNA-seq analysis of the mRNA was performed according to the Chromium NextGem Single Cell 3ʹ Reagent Kits v3.1 user guide. Briefly, a given excess of cells was loaded to the 10x controller to reach a target number of around 1 × 10^4^ a given excess. Quantification of the libraries was performed using Qubit 4 Fluorometer.

ScRNA-seq was conducted using the NextSeq 2000 platform, following the manufacturer‘s protocol. In brief, a single-cell suspension was loaded into the microfluidic chip together with gel beads, reverse transcription reagents, and oil. The microfluidic system partitioned the mixture into thousands to tens of thousands of Gel Beads-in-Emulsion (GEMs), each ideally encapsulating a single cell and a uniquely barcoded bead. Within each GEM, cells were lysed, and mRNA transcripts were captured by the bead-bound barcoded primers. Reverse transcription was subsequently performed inside the GEMs to synthesize complementary cDNA from the captured mRNA. Following reverse transcription, the GEMs were broken to release the barcoded cDNA, which was then amplified by PCR to generate sufficient material for downstream processing. The amplified cDNA was used to construct sequencing libraries compatible with Illumina sequencing platforms through the addition of sequencing adapters. Finally, the prepared libraries were sequenced on Illumina instruments, producing paired-end reads containing both cell barcodes and transcript sequences. The resulting RNA sequence data were aligned to the human reference genome for further analysis.

### ScRNA-seq Raw data processing

The 10x Genomics Cell Ranger analysis pipeline (version 7.0.0) was employed using default parameters. Briefly, binary base call (BCL) files were demultiplexed into FASTQ files using the cellranger mkfastq command, incorporating the appropriate sample sheet and 10x Genomics-supplied barcodes. Subsequently, the cellranger count pipeline was used to align sequencing reads to the designated 10x Genomics reference genome, quantify reads aligning to each gene (including intronic reads in the count matrix), and perform initial clustering and generate summary statistics. Finally, the outputs from the cellranger count analyses for all samples were aggregated and normalized to equivalent sequencing depths, and feature- barcode matrices were recomputed using the cellranger aggr function.

### RNA velocity analysis

To infer differentiation trajectories of MES, CPCs, and DE cell clusters, we employed velocyto (v0.17.17) to quantify spliced and unspliced RNA reads using default parameters. Subsequent analyses and visualizations were performed with the Python packages scVelo (v0.2.4) and Dynamo (v1.1.0) for each dataset. Marker gene expression across different cell clusters was visualized using violin plots generated by the VlnPlot function from the Seurat package. These violin plots illustrate the distribution of marker gene expression levels within the respective cell populations.

### Bulk RNA-seq library prep

For bulk RNA-seq libraries, we extracted RNA from the WT and *PTEN^−/−^* MES cells and CPC cells using TRIzol (Invitrogen) according to the manufacturer‘s protocol. Bulk RNA-seq libraries were prepared by Ouyi Biomedical Technology Co., Ltd (China) using the NEBNext Ultra II RNA Library Prep Kit (New England Biolabs, E7805S). The resulting libraries exhibited fragment sizes ranging from 220 to 500 bp, with a peak at approximately 350 bp. Sequencing was performed on an BGISEQ-500 platform, generating 150-bp paired-end reads. Differentially expressed genes (DEGs) were identified based on the criteria of expression percentage > 0.1, log2 fold change > 0.25, and adjusted p-values. Volcano plots were generated using the ―geom_point‖ function in the R package ggplot2. Principal component analysis (PCA) was conducted using the getDimRedPlot function from the muRtools R package (v0.9.5), utilizing the first two principal components.

### Bulk ATAC-seq

Bulk ATAC-seq libraries were prepared according to the ATAC-Seq Kit Manual (Active Motif, # 53150). Briefly, 50,000 to 100,000 fresh cells were collected, and nuclei were extracted using ATAC lysis buffer. Subsequently, assembled Tn5 transposomes and 2 × Tagmentation Buffer were added to the nuclei, followed by incubation to simultaneously fragment the DNA and insert sequencing adapters (tagmentation). The tagmented DNA was then purified using the provided DNA purification columns and buffers. PCR amplification was performed using Q5 High-Fidelity DNA Polymerase and indexed primers (i5 and i7) to generate Illumina-compatible libraries. Amplified libraries were cleaned with SPRI beads to remove primers and small fragments. Library size distribution was assessed using a fragment analyzer, with an expected size range of approximately 250– 1000 bp and a characteristic ∼150 bp periodicity reflecting nucleosome spacing. Libraries were sequenced using paired-end illumina sequencing, targeting a depth of at least 20–30 million reads per sample.

The ATAC-seq signal for each gene was quantified by summing the signal values of all peaks located within the gene. Peak signal values were calculated using Generich. Bubble plots were generated to display both the ATAC-seq signal and RPKM values for selected transcription factors. The difference in ATAC-seq signal between day 3 and day 6 was calculated as the mean ATAC-seq value of samples collected at day 6 minus the mean value of samples collected at day 3.

### The wordcloud figure method

The ATAC-seq signal for each gene was quantified by summing the peak signal intensities located within the gene region. Peak signal values were calculated using Generich. The differential ATAC value between day 3 and day 6 was determined by subtracting the mean ATAC value at day 3 from the mean value at day 6 across samples. A word cloud was generated based on these data, where the frequency of transcription factors was replaced by their corresponding differential ATAC values between day 3 and day 6. The word cloud visualization was created using the ‗wordcloud2‘ R package, with larger characters representing transcription factors exhibiting greater differences in ATAC accessibility.

### Comparison of single-cell transcriptomic dataset among MES organoids and monkey embryo

Natural primate embryo scRNA-seq datasets utilized for integration analysis were obtained from previously published studies^27^. Human MES organoid scRNA-seq data were projected onto the monkey embryo UMAP embedding to facilitate comparative analysis. Quality control and filtering of cells were performed using the Seurat v4.1.1 package, followed by data normalization. The scRNA-seq data for the monkey embryo were retrieved from the NCBI Gene Expression Omnibus (GEO) under accession number GSE193007. The provided count expression matrix and cell annotation files were used to construct a Seurat object with the Seurat (v4.1.1) R package. The monkey embryo reference dataset comprised 26,135 genes and 56,636 cells, with cell identities annotated according to the metadata file MEF56636-meta.csv.The human MES organoid and monkey embryo Seurat objects were merged using the ‘merge’ function and subsequently integrated via the ‘IntegrateLayers’ function. The merged dataset was normalized using ‘NormalizeData’ and scaled with‗ScaleData‘. Uniform Manifold Approximation and Projection (UMAP) embeddings were computed using the ‘RunUMAP’ function. Cell annotations from the MES organoid Seurat clusters were appended to the merged object, labeling these cells accordingly, while monkey embryo cells were annotated as ‘monkey embryo’. Additionally, annotations from the reference dataset were incorporated, with MES organoid cells labeled as ‘APS organ’. Both annotations were visualized on the UMAP plots to distinguish cell populations.

### Histological Analysis

Mouse embryos were fixed overnight in 4% PFA, embedded in paraffin, and sectioned at a thickness of 5 µm. For immunohistochemistry, tissue sections were blocked for 30 minutes at room temperature, followed by incubation with primary antibodies overnight at 4°C. The anti-FoxA2 antibody was employed for immunofluorescence analyses.

### Phosphoproteomic analysis

In this study, each sample was independently prepared, with proteins enzymatically digested and phosphorylated peptides subsequently enriched prior to analysis by data-independent acquisition (DIA) mass spectrometry. The resulting DIA raw data files were imported into Spectronaut software for comprehensive analysis. For each sample, the numbers of identified phosphorylation sites, modified peptides, and phosphorylated proteins were quantified. To assess the distribution of phosphorylation sites across proteins, the total number of Class I phosphorylation sites detected on all identified proteins was enumerated. Among the modified proteins, the average frequency of Class I phosphorylation sites was calculated as 0.8 sites per 100 amino acids.

To identify differentially expressed phosphorylated proteins between experimental groups, the dataset was further filtered to retain statistically significant modifications. Recognizing that individual proteins may harbor multiple phosphorylation sites exhibiting distinct regulatory patterns, quantification was performed at the modified peptide level rather than at the protein level. Differentially modified peptides were defined using a fold-change threshold of ≥1.5 for upregulation or ≤0.67 for downregulation, combined with a significance cutoff of p < 0.05 (Student‘s t-test or equivalent). This approach enabled the identification and quantification of significantly upregulated and downregulated phosphorylated peptides between comparison groups.

### Statistics and reproducibility

At least three independent experimental replicates were performed for each condition, with each replicate conducted in triplicate. Statistical analyses were carried out using GraphPad Prism software (version 6). Data are presented as mean ± standard deviation (SD). The sample size (n) corresponds to the number of independent biological replicates. Statistical significance between groups was assessed using an unpaired Student‘s *t*-test. A threshold of *p* < 0.05 was considered statistically significant. Significance levels are denoted as follows: ns, not significant (*p* > 0.05); \**p* < 0.05; \*\**p* < 0.01; and \*\*\**p* < 0.001. RNA-seq, ATAC-seq, phosphoproteomic, and metabolism experiments were performed in biological duplicate.

## Data availability

The high-throughput sequencing datasets have been deposited in the Gene Expression Omnibus (GEO) database. The accession numbers are GSE301515 for single-cell RNA-seq, GSE301159 for bulk RNA-seq, and GSE301161 for ATAC-seq. Additionally, the metabolism datasets have been submitted to the NIH Metabolomics Workbench under the accession number datatrack_id:6182. The phosphoproteomics datasets have been deposited in the PRIDE repository, with the accession number PXD065688.

## Supporting information

Supplementary Video 1 -MES organoid

Supplementary Video 2 - EC organoid

## Acknowledgements

This work was supported by a Natural Science Foundation of China (NSFC)-Young Scientists Fund (#82200314), Innovation and Technology Commission of the Hong Kong Government grants (GHX09721SZ and MHP00523), Direct Grant for Research (#2021.069 and #2022.082) from CUHK, the National Natural Science Foundation of China (NSFC)/Research Grants Council (RGC) Joint Research Scheme (#N_CUHK428/22), and Strategic Seed Funding for Collaborative Research Scheme (#MK/WW/SSFCRS2324/8014/24jh) from CUHK, the State Ministries Special Budget to support the MOE Key Laboratory for Regenerative Medicine (CUHK–Jinan University) [2622009], partially supported by Science and Technology Planning Project of Guangdong Province, China (2023B1212120009), CUHK Laboratory Support Special Fund for Key Laboratory for Regenerative Medicine, Ministry of Education, China, and Shenzhen Virtue University Park Laboratory Support Special Fund (YFJGJS1.0), and the Chinese University of Hong Kong (CUHK)–Shandong University (SDU) Joint Laboratory on Reproductive Genetics. This work was also partially supported by Center for Neuromusculoskeletal Restorative Medicine Limited, Hong Kong.

## Author Contributions

W.W. performed the majority of the experiments. Experimental design was developed by W.W., H.L., G.L., and W.-Y.C. Cell culture and molecular assays were conducted by Q.H. and H.Z., while bioinformatics support was provided by S.-W.C. and Z.X. Cryosectioning and staining of organoids were carried out by H.T., and animal experiments were performed by Y.L., X.S., and Y.G. Data analysis was facilitated by Y.F., X.L., X.B., and A.M.C. The study was conceived by H.L., G.L., and W.-Y.C. Data interpretation and manuscript preparation were undertaken by W.W., H.L., G.L., and W.-Y.C., with contributions from all other authors.

