## Extended Figures for "Human organoids reveal PTEN-driven mesendoderm specification via retinoic acid signaling suppression"

**Extended Data Figure 1. MES organoids generation from hiPSCs.**
***a.*** MES organoids were harvested at multiple time points and subjected to IF staining with anti-SOX17 and anti-EOMES antibodies. Scale bar, 100 μm. ***b***. Temporal expression dynamics of several MES marker genes were quantitatively analyzed. ***c***. Whole-mount IF staining of MES organoids was performed to assess spatial marker distribution. Scale bar, 100 μm. ***d***. Whole-mount IF staining of EC organoids was conducted using an anti-CDH5 antibody, accompanied by phase-contrast and bright-field imaging. Scale bar, 200 μm. ***e***. 3D reconstructions of EC organoids was achieved via fluorescence microscope, Scale bar, 200 μm. Single cells dissociated from EC organoids are shown. Scale bar, 100 μm.

**Extended Data Figure 2. scRNA-seq analysis of MES organoids.**
***a***. and ***b***. Identification of APS, Epi, and CPC cell clusters, illustrated with corresponding violin plots depicting gene expression distributions.

**Extended Data Figure 3. *PTEN* ablation impairs MES formation and DE lineage specification.** ***a***. Analysis of MES organoids derived from WT and *PTEN*$^{-/-}$ groups. Scale bar, 100 μm. ***b***. Single cells isolated from WT and *PTEN*$^{-/-}$ MES organoids were replated for further analysis. Scale bar, 100 μm. ***c***. IF staining of iPSCs-derived DE cells co-expressing *SOX17 and FOXA2*. Scale bar, 50 μm. ***d***. Quantification revealed a significant reduction in *SOX17- and FOXA2*-positive cells following *PTEN* ablation. ***e***. Expression levels of DE markers (*FOXA2*, *SOX17*) were decreased in *PTEN*$^{-/-}$ DE cells. ***f.*** Flow cytometry analysis demonstrated a marked decrease in the proportion of DE cells upon *PTEN* deletion. ***g***. GO enrichment analysis identified key signaling pathways implicated in DE differentiation. Statistical test, two-sided *t*-test; $*P < 0.05$; $**P < 0.01$; $***P < 0.001$.

**Extended Data Figure 4. *PTEN* ablation suppresses cardiac mesoderm formation.**
***a***. IF staining showed a significant reduction in cardiac progenitor cells following *PTEN* ablation. Scale bar, 100 μm. ***b***. Flow cytometry analysis demonstrated a marked decreased in the proportions of GATA4-positive and NKX2.5-positive cells in *PTEN*$^{-/-}$ cells. ***c***. Generation of hCO from hiPSCs was performed. ***d***. IF staining confirmed expression of the CPC marker NKX2.5 in both WT and *PTEN*$^{-/-}$ hCOs. Scale bar, 100 μm. ***e***. Expression levels of cardiac mesoderm markers were significantly diminished in *PTEN*$^{-/-}$ hCOs. Statistical test, two-sided *t*-test; $*P < 0.05$; $**P < 0.01$; $***P < 0.001$.

**Extended Data Figure 5. EC lineage specification is impaired by *PTEN* ablation.**
***a***. Phosphoproteomic profiling reveals disruption of the VEGFA–VEGFR2 signaling pathway in *PTEN*$^{-/-}$ cells.

**Extended Data Figure 6. *PTEN*-OE rescues MES cell generation and convergent migration.**
***a***, ***b***. *PTEN*-OE reinstated MES formation, as evidenced by the co-expression of SOX17 and EOMES. Scale bar, 100 μm. ***c***. The mRNA levels of MES marker genes, including *EOMES, GSC, FOXA2, LHX1*, and *SOX17,* were significantly restored in the *PTEN*-OE group. ***d***. Convergent migration of MES was markedly rescued following *PTEN*-OE. Scale bar, 50 μm. ***e***. ATAC-seq reveals the reduced chromatin accessibility at MES marker loci (*GSC*, *LHX1, FOXA2*) at day 3. Statistical test, two-sided *t*-test with Benjamini–Hochberg correction; $*P < 0.05$; $***P < 0.001$.

**Extended Data Figure 7. PTEN modulates MES cell generation via its phosphatase activity.** ***a***. Western blot analysis demonstrated that the PI3K inhibitor PX-866 effectively restored MES cell generation. ***b***, ***c***. The induction of MES cells was similarly rescued by PX-866 treatment, as evidence by increased MES marker expression. Scale bar, 100 µm. ***d***. Quantitative analysis showed that mRNA levels of MES-specific marker were significantly restored following PX-866 treatment. Statistical test, two-sided *t*-test with Benjamini–Hochberg correction; *$P < 0.05$; ***$P < 0.001$.

**Extended Data Figure 8. PTEN regulates MES cell generation via CYP26A1-mediated RA signalling.** ***a***. Knockdown efficiency of CYP26A1 was validated by Western blot analysis, demonstrating reduced protein levels following shRNA treatment. ***b***, ***c***. *CYP26A1* knockdown significantly impaired MES cell generation. Scale bar, 100 µm. ***d***, Sanger sequencing for two *CYP26A1* deleted iPSC colonies generated by CRISPR/Cas9 system. ***e***. *CYP26A1* ablation significantly inhibited MES cell generation. Scale bar, 100 µm. ***f***. Expression levels of MES marker genes were significantly decreased in $PTEN^{-/-}$ MES cells. Statistical test, two-sided *t*-test; *$P < 0.05$; **$P < 0.01$; ***$P < 0.001$.

**Extended Data Figure 9. WNT/β-catenin signalling is implicated in the PTEN-mediated regulation of MES generation.** ***a***. Western blot analysis demonstrated that treatment with PI3K inhibitor PX-866 significantly restored the protein level of CYP26A1 and β-catenin. ***b***. PX-866 treatment markedly increased the mRNA level of CYP26A1 in $PTEN^{-/-}$ MES cells. ***c***. *PTEN*-OE significantly rescued CYP26A1 expression. ***d***. Kinome dendrogram derived from human phosphoproteomics data revealed enrichment of WNT/β-catenin signaling when comparing WT and $PTEN^{-/-}$ MES cells. ***e***. Heatmap analysis indicated PTEN-dependent regulation and enrichment of WNT signaling components. ***f***. The diagram illustrates that PTEN may regulate CYP26A1 expression through the WNT/β-catenin signaling pathway. Statistical test, two-sided *t*-test with Benjamini–Hochberg correction; *$P < 0.05$; ***$P < 0.001$.

**Extended Data Figure 10. Excessive RA exerts a detrimental effect on MES cell generation.** ***a***. Excessive RA significantly suppressed MES generation, mirroring the phenotype observed in the $PTEN^{-/-}$ group. ***b***. Treatment with increasing concentrations of RA resulted in a dose-dependent inhibition of MES cell generation. Scale bar, 100 µm. ***c***. Excessive RA exposure reduced the proportion of MES cells within MES organoids. Scale bar, 100 µm. ***d***, ***e***. Administration of an RA antagonist enhanced MES cell generation, as evidenced by increased numbers of EOMES- and SOX17-positive cells and elevated mRNA levels of MES markers. Scale bar, 100 µm. ***f***. The RA antagonist significantly promoted MES organoid formation relative to the DMSO-treated group. Statistical test, two-sided *t*-test; *$P < 0.05$; **$P < 0.01$; ***$P < 0.001$.

### Extended Data Figure 1

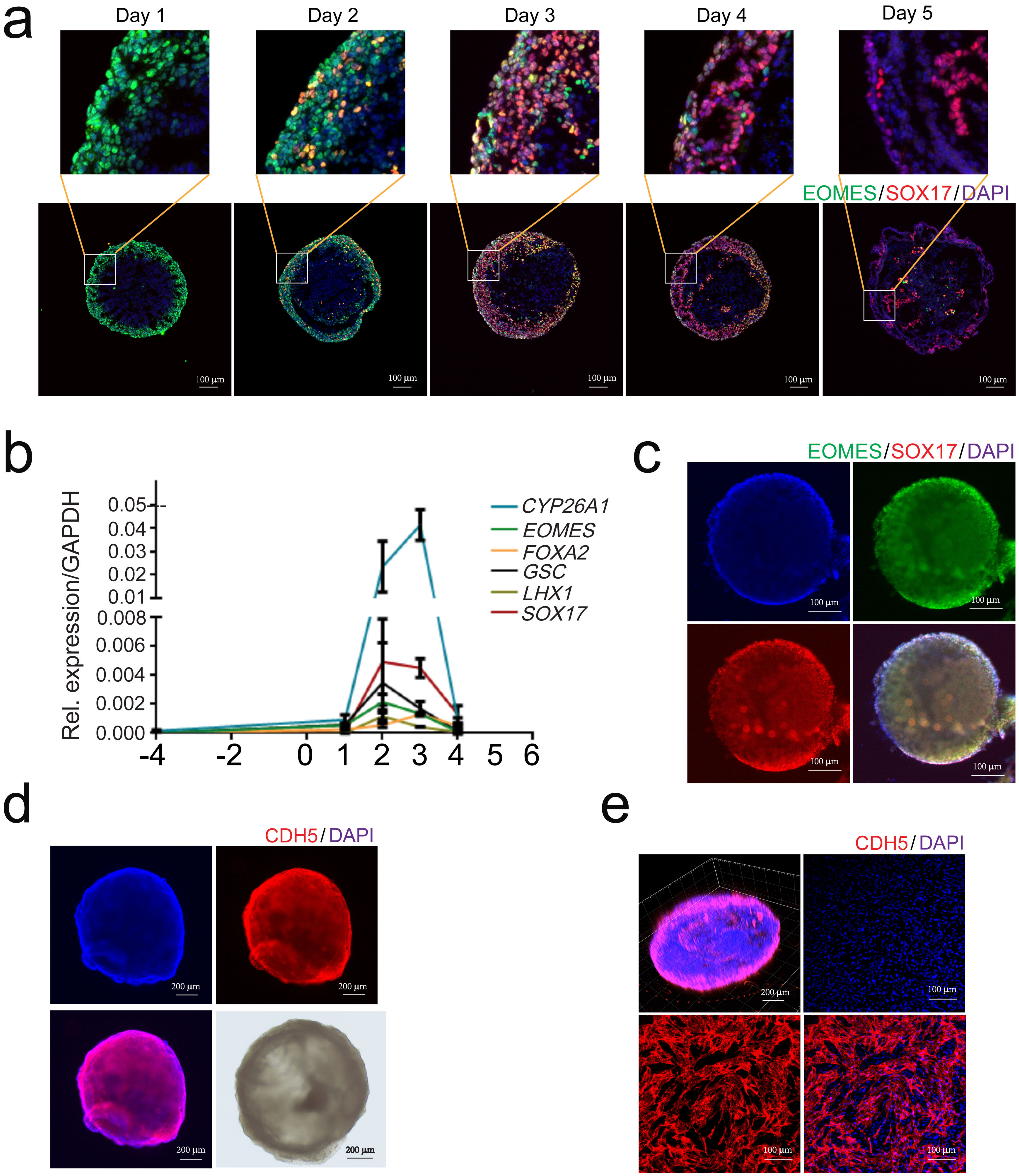

### Extended Data Figure 2

a

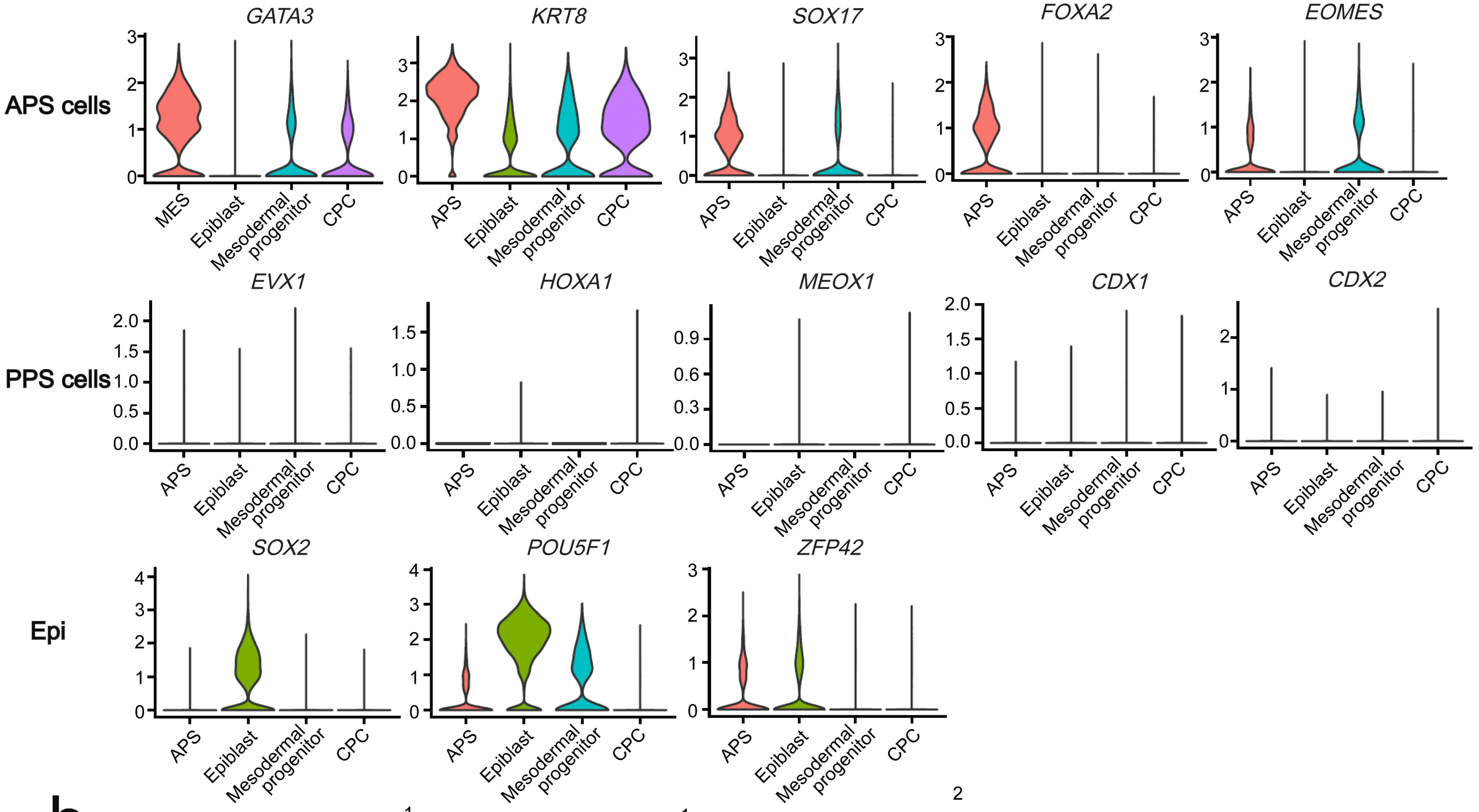


b

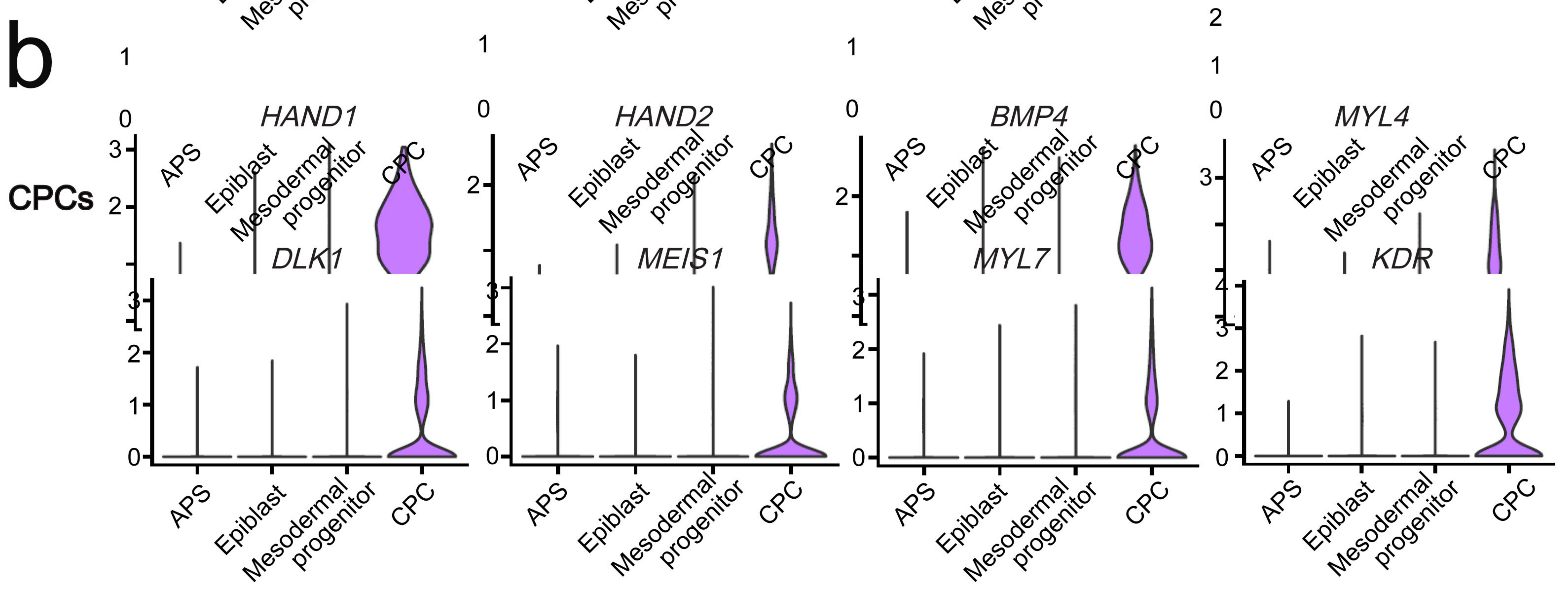

### Extended Data Figure 3

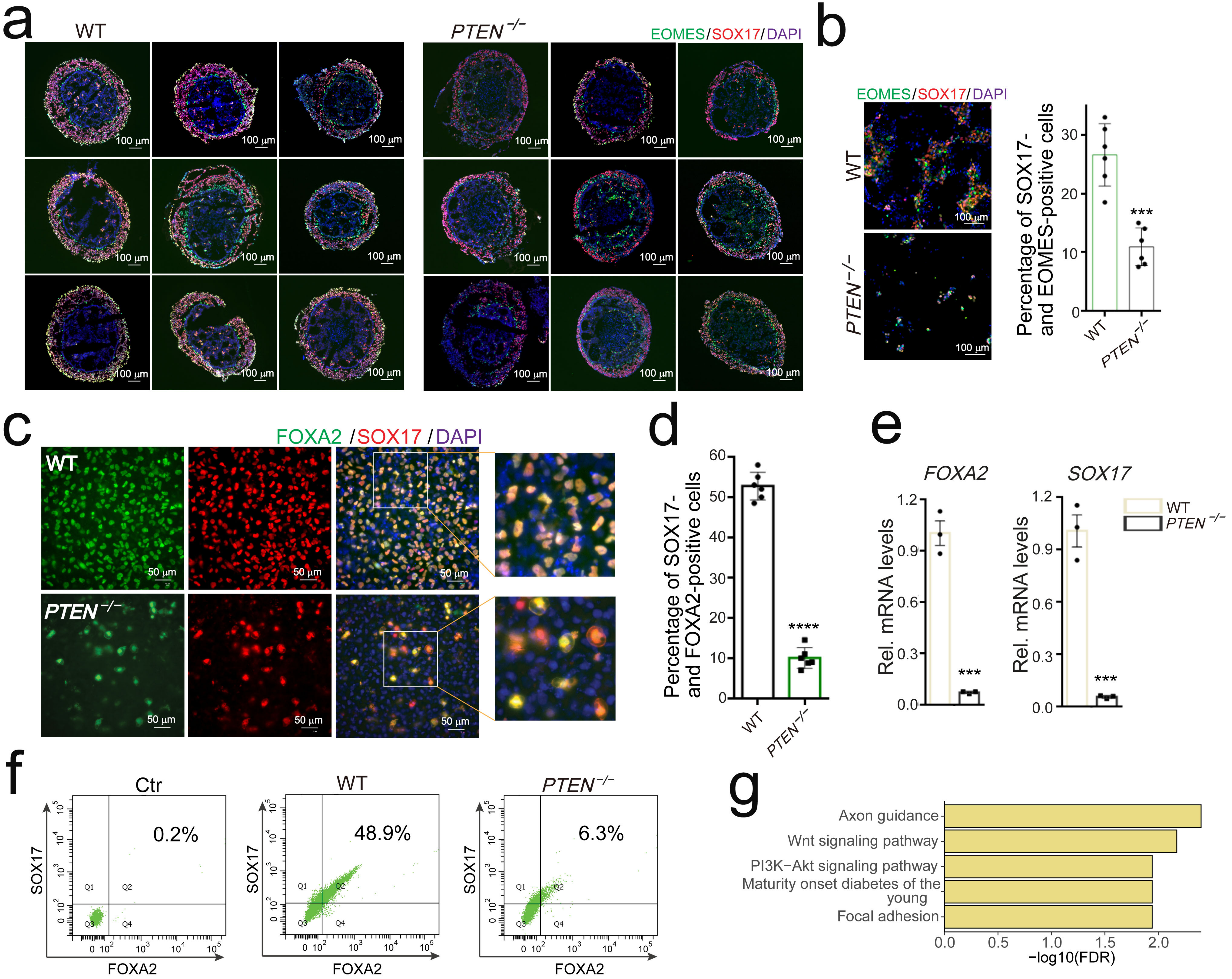

### Extended Data Figure 4

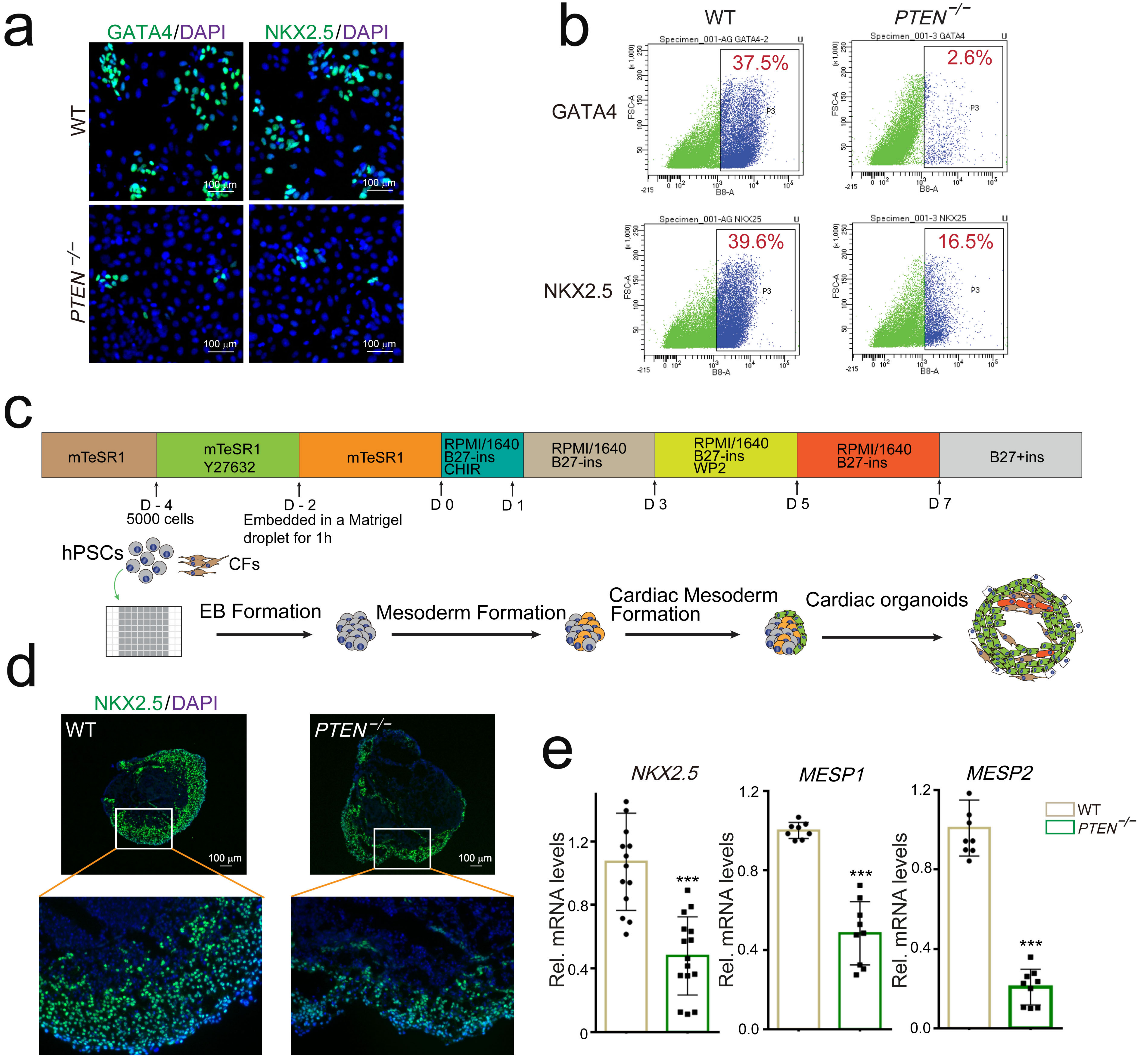

### Extended Data Figure 5

a

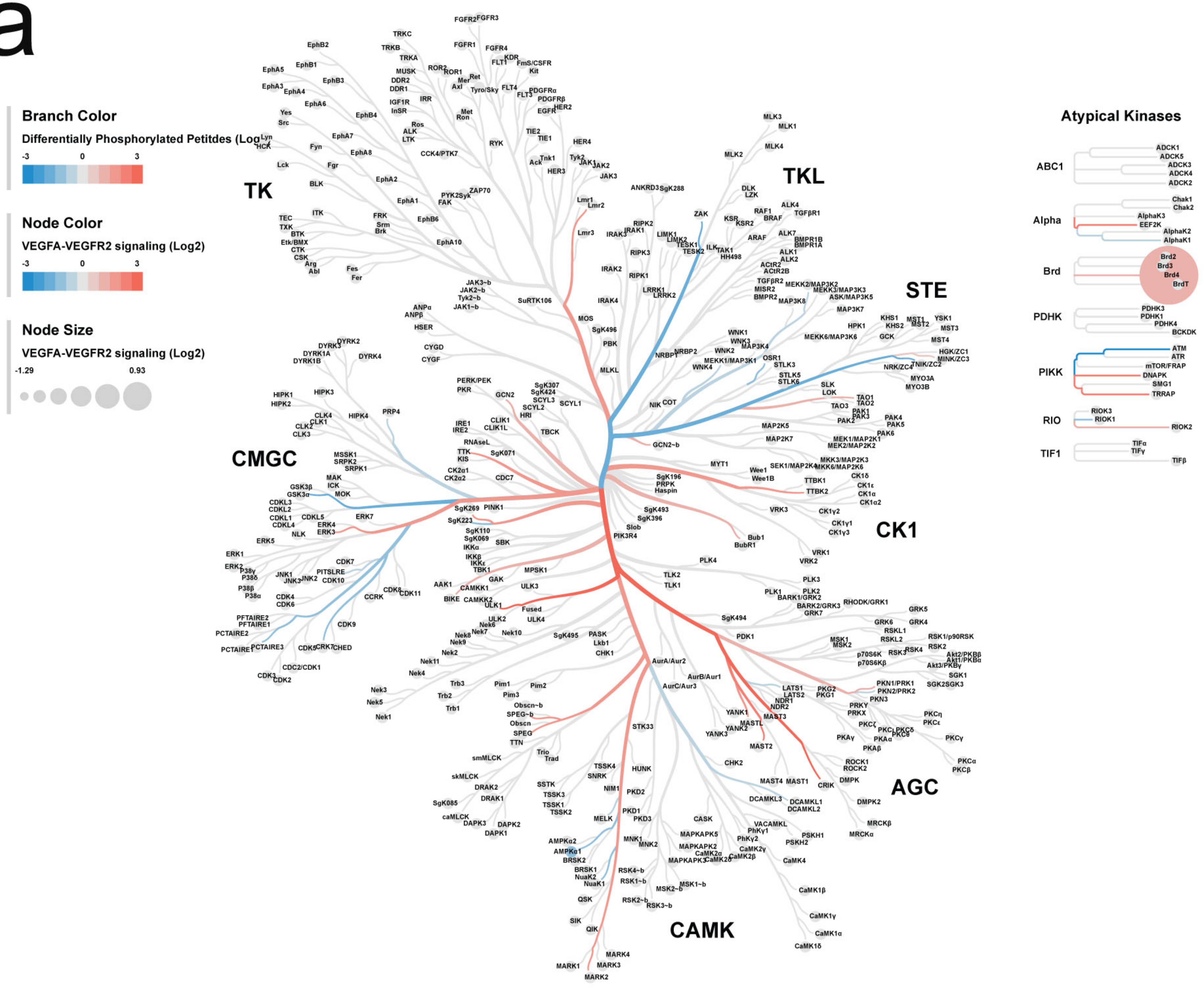

### Extended Data Figure 6

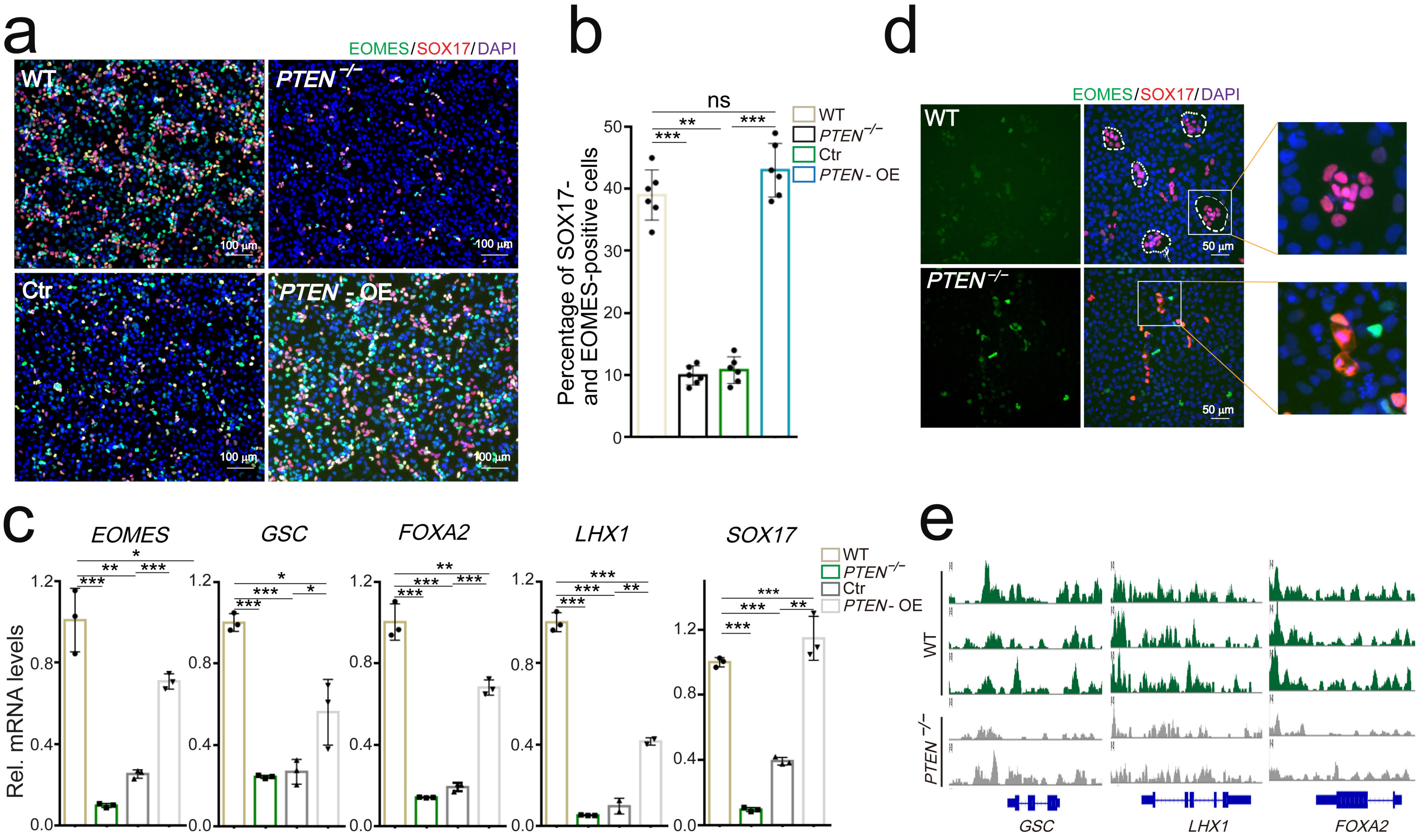

### Extended Data Figure 7

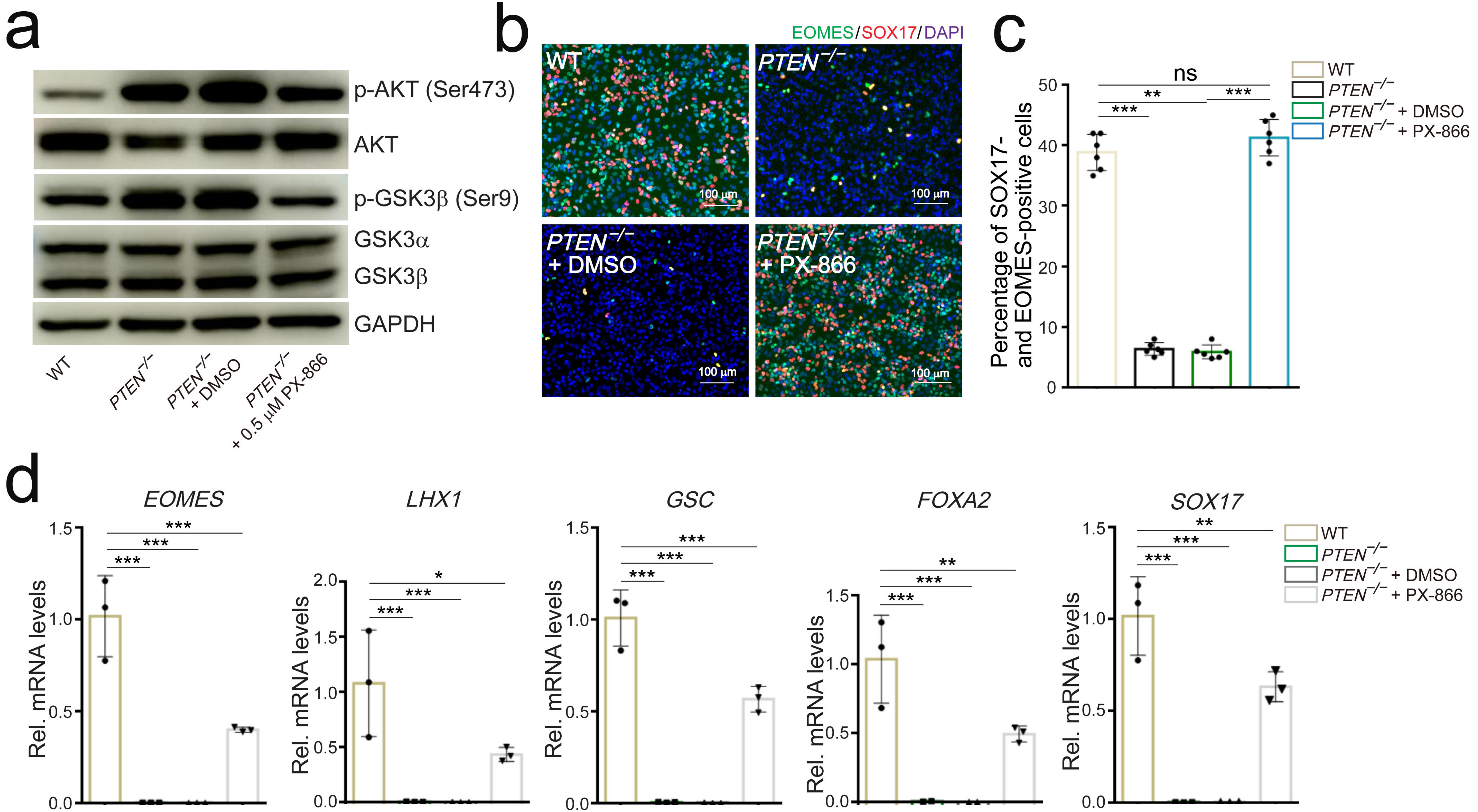

### Extended Data Figure 8

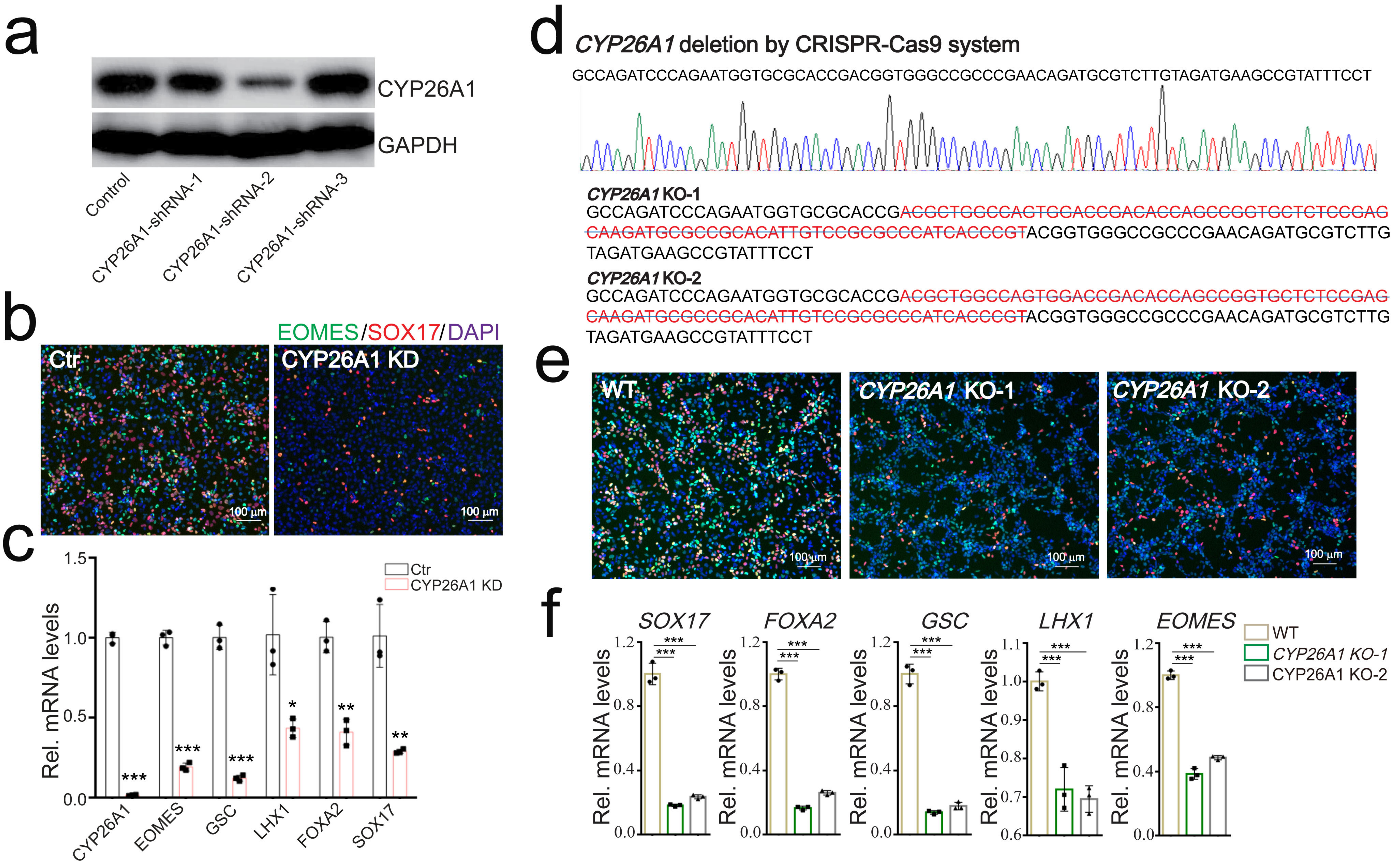

### Extended Data Figure 9

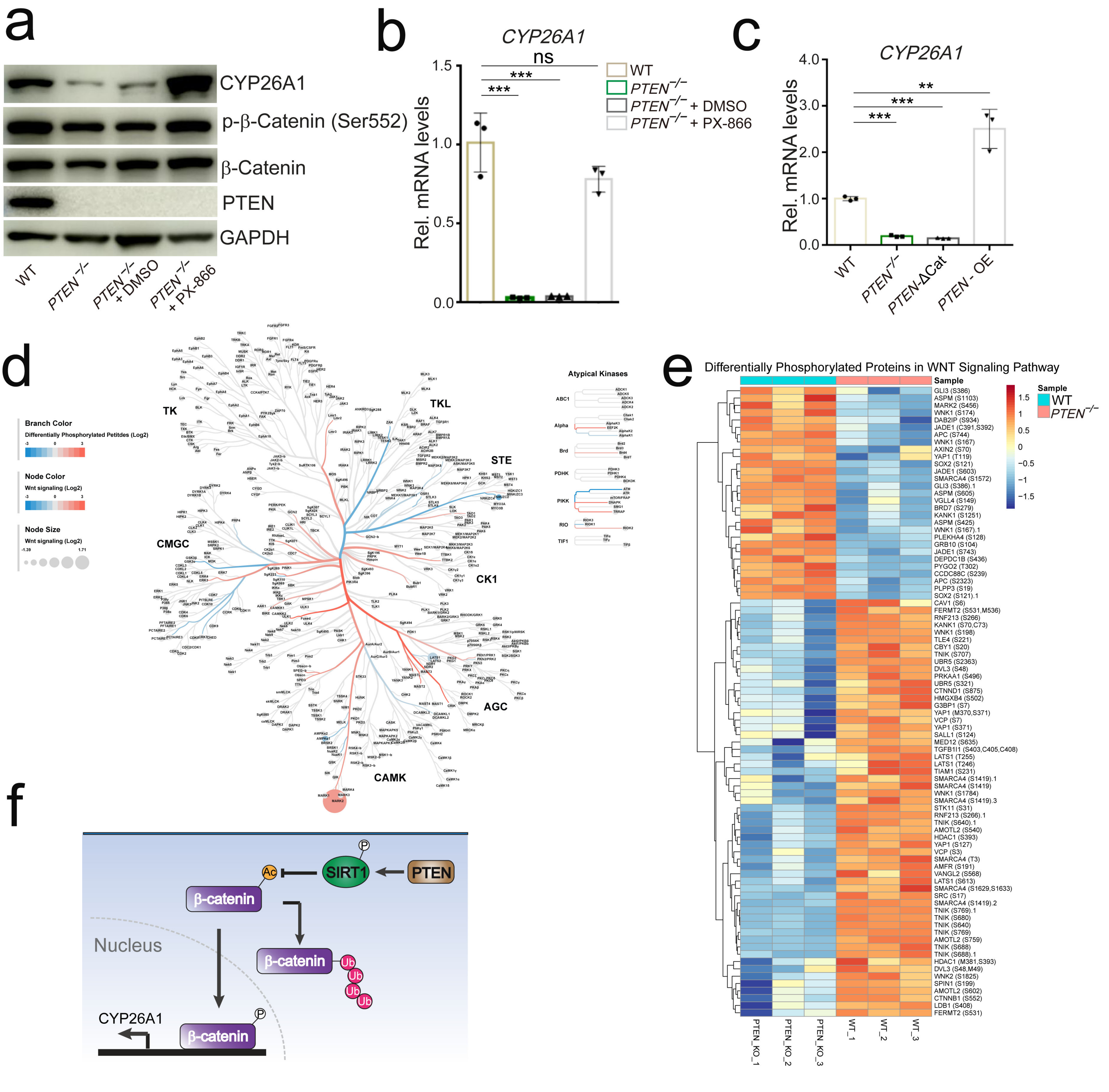

### Extended Data Figure 10

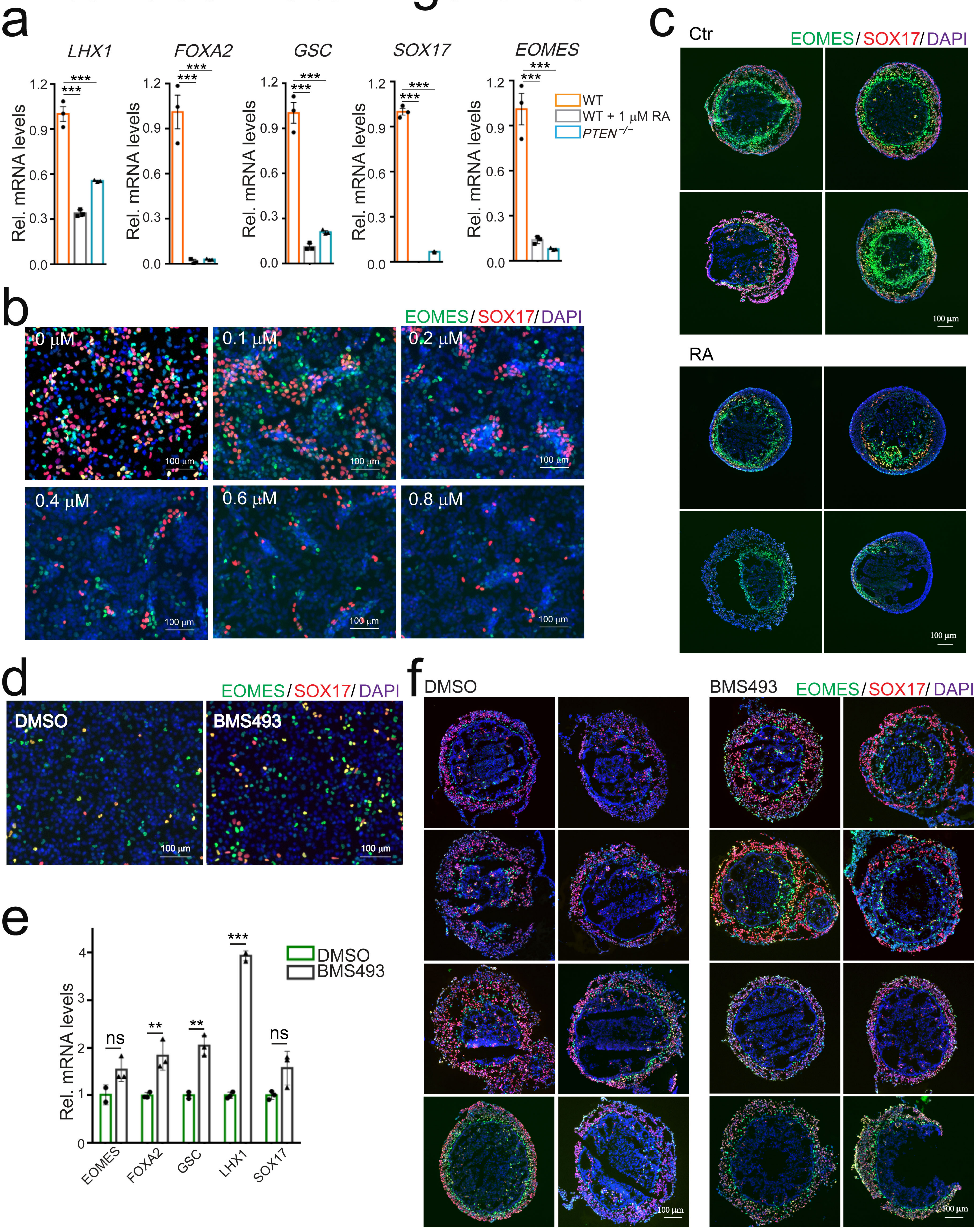
